# From cell size and first principles to structure and function of unicellular plankton communities

**DOI:** 10.1101/2022.05.16.492092

**Authors:** K.H. Andersen, A.W. Visser

## Abstract

Here we review, synthesize, and analyse the size-based approach to model unicellular plankton cells and communities. We first review how cell size influences processes of the individual the cell: uptake of dissolved nutrients and dissolved organic carbon, phototrophy, phagotrophy, and metabolism. We parameterise processes primarily from first principles, using a synthesis of existing data only when needed, and show how these processes determine minimum and maximum cell size and limiting resource concentrations. The cell level processes scale directly up to the structure and function of the entire unicellular plankton ecosystem, from heterotrophic bacteria to zooplankton. The structure is described by the Sheldon size spectrum and by the emergent trophic strategies. We develop an analytical approximate solution of the biomass size spectrum and show how the trophic strategies of osmotrophy, light- and nutrient-limited phototrophy, mixotrophy, phagotrophy depend on the resource environment. We further develop expressions to quantify the functions of the plankton community: production, respiration and losses, and carbon available to production of higher trophic levels, and show how the plankton community responds to changes in temperature and grazing from higher trophic levels. We finally discuss strengths and limitations of size-based representations and models of plankton communities and which additional trait axes will improve the representation of plankton functional diversity.

## 1 Introduction

The pace of global change spurs the imperative for predictive, global scale models of marine ecosystems. Key questions that confront us are how the diversity and functioning of marine ecosystems will change, how these changes will impact key ecosystem services such as primary and secondary production, ocean oxygen concentration and carbon sequestration, and whether these services are subject to tipping processes. Traditional models, that have been lovingly calibrated and validated to current-day situations, and through which we have learned so much of marine ecosystem dynamics, are challenged with this task. The world is moving rapidly out of the calibration envelopes for which they were calibrated, and the validation of model predictions with observed ecosystems can no longer be the sole gold-standard measure of model success. In an ideal world predictive, global scale models should be rooted in “first principles”: the rules of the natural world whose validity are considered fundamental and unchanging. In this context, mass and energy conservation, chemical reaction kinetics and evolution by natural selection can be considered examples of first principles. Models of ecosystems do not have recourse to such first principles *per se*. Nevertheless, individual organisms are constrained by first principles that are manifested at all scales of life, from the reaction kinetics and topology of life’s fundamental molecules, the physical limitations of functions of the cells, the circulatory systems, and the geometry of the body plan. One aspect of life where first principle constraints are most evident is in relations to the size of individual organisms (Haldane, 1926; Andersen et al., 2016). Here we attempt to scale from individual organism to ecosystem structure and function. We use unicellular planktonic life as an example where first principles constraints on the individual cell have a particularly strong effect on the ecosystem structure and function (Kiørboe, 1993).

Unicellular plankton is an incredibly diverse group of organisms. Taxonomically they represent four domains of life: archaea, bacteria, algae and protozoa. In terms of cell size, plankton spans 8 orders of magnitude in mass, the same range as between a beetle and an elephant. Functionally, unicellular planktonic ecosystems show the entire range of trophic strategies of primary producers (phytoplankton), grazers and predators (zooplankton), and detritivores (bacteria). The unicellular planktonic food web drives the fast turnover of inorganic dissolved matter in the oceans, half the global primary production, the main carbon flux from the photic zone, and the turn-over of inorganic and organic matter in the world’s oceans and lakes. All metazoans – multicellular plankton, jellies, fish, benthic organisms, and marine mammals – rely on surplus production from unicellular plankton food webs (Ryther, 1969; Stock et al., 2017). Without the unicellular food web, macroscopic life in the oceans would be extremely impoverished.

The difficulty of observing and experimenting with unicellular plankton food webs have put models in a central position, not just for predictions of responses to changes, but also for understanding the structure and function of the ecosystems. Any ecosystem model faces a choice of how to represent the diversity of organisms. The classic food web approach, which is often applied to higher trophic levels, attempts to resolve all populations and their interactions with other populations. This approach only works for smaller isolated ecosystems and is clearly unsuited to unicellular plankton where we rarely have a clear overview of the full taxonomic diversity. Plankton models instead describe diversity by lumping species into distinct functional groups. The simplest grouping is between phytoplankton (P) and zooplankton (Z), with phytoplankton representing all phototrophic organisms and zooplankton their grazers (Franks, 2002). This grouping together with nutrients (N) lead to “NPZ” models, which have been remarkably successfully in capturing the main features of seasonal succession (Evans and Parslow, 1985; Fasham et al., 1990; Anderson et al., 2015) and global patterns of production (e.g. Palmer and Totterdell, 2001). However, their success is contingent on model parameters being tuned to the observations themselves. In this way, parameters of each group are adjusted to represent the physiology and ecology of the dominant species in the group within the geographic region that is modelled. When conditions change though, other species with different parameters may become dominant and the model no longer represents the new ecosystem (Franks, 2009). This parameter tuning therefore reduces our confidence in the model’s ability to reproduce ecosystem dynamics when conditions change outside the model’s tuning envelope.

A further elaboration of plankton diversity is achieved by breaking the trophic groups into additional functional groups (Anderson, 2005; Le Quere et al., 2005; Hood et al., 2006). The functional groups are often aligned with dominant taxonomic groups including coccolithophores, dinoflagellates, ciliates, and diatoms, or more general groups, e.g., silicifiers, calcifiers etc.. While the functional-group approach introduces additional flexibility and accuracy it does so at the price of increased complexity and additional parameters. Nevertheless, each group still represents a huge diversity of organisms – for example, the size range of diatoms spans from a few tens to 10^7^ cubic micrometers – and parameters for each group are still tuned to represent the dominant species in the modelled region. While the introduction of further realism improves the models fit to observations it does not solve the fundamental problem of parameter tuning. Further, the addition of new functional groups leads into a complexity trap with a proliferation of state variables and parameters.

Size-based models break free of the complexity trap of functional groups by representing the plankton community with size groups that each represent all cells in a given size range regardless of their taxonomic affiliation. Technically, each size group is modelled largely in the same way as a functional groups. The main difference is that the parameters are not independently determined for each size group. Instead, parameters follow from a smaller set of scaling coefficients and exponents that apply to all sizes. In this manner size models are flexible with respect to the number of state variable while retaining a small set of parameters that is, at least in theory, generally valid. Breaking free of the complexity trap in this manner comes at the cost of a poor representation of taxonomic diversity. However, the size-based model provides a framework where functional diversity is an emergent property of the model rather than a consequence of its structure.

There are other reasons for using cell size as the governing axis of diversity. It is now well documented that within plankton many of the fundamental rates and processes scale with cell size (Fenchel, 1987; Kiørboe, 1993; Finkel et al., 2010; Marañón, 2015): affinities for nutrients (Edwards et al., 2012) or light (Taguchi, 1976; Edwards et al., 2015), maximum bio-synthesis rates and respiration rates (Kiørboe and Hirst, 2014), clearance rates (Kiørboe and Hirst, 2014), predator-prey mass ratios (Hansen et al., 1994) and predation risk from larger organisms (Hirst and Kiørboe, 2002). Importantly, many of these scaling relations emerge from fundamental physical limitations due to geometry (light affinity), diffusion (affinity for dissolved organic matter), and fluid mechanics (e.g. Stokes’ law or feeding mechanics (Nielsen et al., 2017)). In other words: the parameters are constrained by first principles from geometry or classical physics. A further advantage of size-based models is the conceptual simplicity that comes from being based on a general description of a single cell. The simplicity extends to the implementation, which only needs a small parameter set and have simpler code. These advantages make size-based descriptions appealing to add diversity within a functional group (Terseleer et al., 2014; Stock et al., 2014) or for the full model structure. Existing size-based models mostly rely on empirical relationships between size and parameters such as half-saturation coefficients, maximum growth rates etc.. This approach facilitates a good fit with observations. Here, we instead try to establish the fundamental mechanisms and strive to determine parameters from fundamental principles by reviewing the literature on the theory of size-based relations with cell size.

Size-based models of plankton have a long history (Armstrong, 1994; Moloney and Field, 1989; Baird and Suthers, 2007; Stock et al., 2008; Banas, 2011; Negrete-García et al., 2022). Size-based concepts are now increasingly used in biogeochemical models to increase the diversity within functional groups according to size (Terseleer et al., 2014; Dutkiewicz et al., 2020; Stock et al., 2014). Most size-based models retain the distinction of functional trophic groups by operating with separate phyto- and zooplankton size distributions (Poulin and Franks, 2010; Ward et al., 2018). A recent strand is purely size-based models where the only difference between cells are their size and no *a priori* distinction between trophic strategy is imposed (Ward and Follows, 2016; Ho et al., 2020; Chakraborty et al., 2020). Such models completely forgo taxonomic-oriented assumptions about the function of the modelled groups (Andersen et al., 2015). All functional differences between size groups and of the community are emergent properties of the model.

There exists an abundance of reviews on the empirical relationships between cell size and various processes (Kiørboe, 1993; Hansen et al., 1994; Finkel et al., 2010; Edwards et al., 2012; Kiørboe and Hirst, 2014; Marañón, 2015; Hillebrand et al., 2021). They tend to focus on phytoplankton and upon describing size-relations as a single power-law function. However, in many cases there is more than one underlying physical process at play. This means that there are transitions between one power-law relation and another, e.g., between nutrient diffusion and surface uptake (Armstrong, 2008) or between maximum synthesis rates and nutrient uptake (Ward et al., 2017). Such transitions at characteristic sizes often lead to important transition in the ecosystem structure (Andersen et al., 2016), for example between phototrophs, mixotrophs, and heterotrophs (Andersen et al., 2015). Identifying characteristic sizes where there is a cross-over between two power-law relations is perhaps even more important for ecosystem structure and function than the power-law relations themselves.

Here we review existing knowledge of size-based relationships for unicellular plankton, from bacteria to zooplankton, and attempt a synthesis that demonstrates the importance of size-based relations for emergent ecosystem structure and function. Our ambition is to identify the first principles responsible for the size-based relations, thereby tying parameters to physical and chemical processes and geometry. Our synthesis show how size-based relationships determine community-level patterns of biodiversity and ecosystem function: the viable size-range, competition, biomass size structure, ecosystem primary and secondary production, and trophic efficiencies. By focusing on the processes related to cell size, we demonstrate the power of these relations for determining community-level patterns and ecosystem functions. The work is organised in five parts. After an initial discussion of the concept of “size” of a cell, we review the relations governing resource uptake, losses, and biosynthesis of a cell, including the theory that links these processes to first principles. Second, we exploit the simple form of the size-based relations to derive analytical solutions for the smallest and largest cell sizes, and for the limiting resources. Third, we scale from the cell-level process to the community size distribution and explore emergent trophic strategies. We derive a scaling solution of the biomass size distribution and explore the trophic strategies and compare with simulations of the size-based model in a chemostat. From the emergent size spectrum and trophic strategies we derive ecosystems functions, including production, and show how the plankton community responds to predation by higher trophic levels or changes in temperature. Our aim is to make a minimal size-based model framework where we prioritize simple conceptual implementation and analytical analysis over capturing complete and accurate biogeochemistry. Nevertheless we show that the model gives reasonable predictions of biomass and production. Overall, our synthesis highlights the importance of fundamental first principles for constraining the unicellular plankton communities and their related functions. We finish by discussing the limitations of the size-based approach and prioritize which additional traits will best improve the representation of functional diversity.

## 2 Measures of cell size

The size of a cell can be measured in two ways: by its physical size – radius or volume – or by its mass, e.g., mass or moles of carbon or nitrogen. There is no universally optimal measure; for some processes physical size is most relevant, for some it is the mass, and for others both measures of size matter. For example, the settling velocity due to Stokes’ law is determined by both the physical size and the mass of the cell. In general, physical size is mostly used to describe limitations due to geometry, e.g., surface limitation, while mass is used to describe metabolism and mass budget of the cell.

Unicellular plankton display an astonishing diversity in cell shape (Ryabov et al., 2021). The functional role of cell shape is largely unknown, though it is conjectured to be related to defence from predation (Smetacek, 2001). For simplicity we ignore the diversity of shapes (except for its relation to the minimum size in Section 4.1), and consider cells to be spherical with physical size characterized by radius *r*. Conversion between physical size (equivalent spherical radius *r*) and mass of substance *X*, *m_X_* is then:

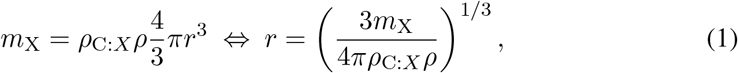

where *ρ* is the carbon density (carbon mass per volume) and *ρ*_C:*X*_ is the elemental mass ratio between carbon and *X*.

In the following, mass is considered as carbon mass and the subscript C is suppressed *m* = *m*_C_. For the theoretical calculations we use a density of *ρ* = 0.4*·*10^−6^ μgC/μm^3^ and Redfield elemental ratios. Conversion between physical size and mass needs to account for differences in density. In particular diatoms are special due to their vacuole which lowers their density. Here we use the comprehensive compilation of Menden-Deuer and Lessard (2000) that explicitly distinguishes between diatoms and other protists to convert observations of cell size to cell mass.

Not all of the cell’s mass *m* is available for functions of biosynthesis (ribosomes), light harvesting (chloroplast) etc. Some part of the cell is devoted to the cell membranes, DNA and RNA, (Kempes et al., 2016). The cell membrane and cell wall takes up a fraction of the cell mass (Raven, 1994; Marañón, 2015). For a spherical cell the fraction of the cell used by the membrane and wall is approximately:

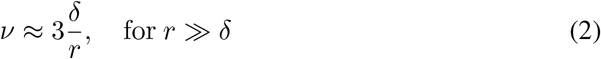

where *δ* ≈ 50 nm is the thickness of the cell wall and membrane (adjusted a bit down from 70-80 nm as given by Raven (1987) to correct for the approximation used in Eq. 2). The effective functional mass is therefore *m*(1 *− ν*). As the cell wall fraction scales as 1/*r*, small cells will be severely limited in the functions due to the material cost of cell membrane and wall. Kempes et al. (2016) further considered the limitation of DNA and RNA, however, the most limiting factor was the cell wall and membrane.

## 3 Effects of cell size on fundamental rates: resource uptake, losses, and biosynthesis

Ecosystem dynamics are driven by individual cells acquiring and processing resources, eventually leading to cell division and cell growth. This section reviews how cell size determines the uptakes of resources: dissolved nutrients, inorganic carbon through photo-harvesting, dissolved organic carbon, and feeding on other, typically smaller, organisms, and how these uptakes are determined by first principles. Some of the acquired resources are lost through passive exudation or used for respiration. The remaining resources are used for biosynthesis (Fig. 1).

**Figure 1:**
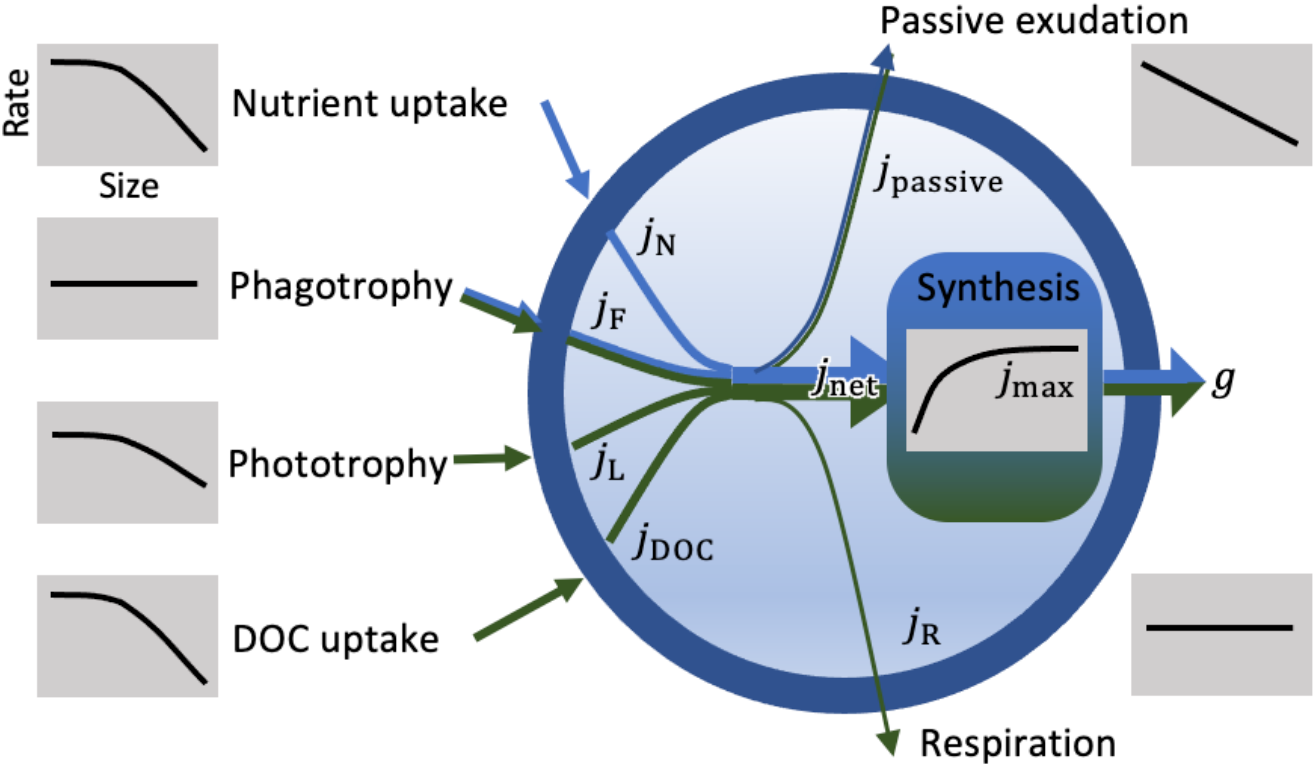
Sketch of the fluxes of nutrients (blue) and carbon (green) in and out of a cell. The grey insets sketches the size-dependency of each mass-specific rate (units of 1/time). Uptakes of nutrients *j*_N_, food *j*_F_, photoharvesting *j*_L_, and dissolved organic carbon *j*_DOC_ are subjected to losses from respiration *j*_R_ and passive exudation *j*_passive_ before they are synthesised with a maximum rate *j*_max_. The end result is the growth (i.e. division) rate *g*.

### 3.1 Resource uptake

Cells take up resources through three mechanisms: diffusive uptake of dissolved organic carbon (DOC) and inorganic matter (*N*), photoharvesting of light (*L*), and phagotrophic uptake of particulate matter (*F*). The potential uptake of resource *X* is proportional to the resource concentration:

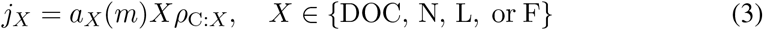

where *j_X_* is the mass specific flux (in units of g_C_/g_C_/time), *a_X_* is the mass specific affinity (volume/day/g_C_), *X* is the resource in units of *X* per volume, and *ρ*_C:*X*_ = *ρ*_C:N_ is the C:N ratio for diffusive uptake of nutrients. By multiplying N uptake with the fixed C:N ratio all the fluxes are measured in the same units and are therefore directly comparable (uptake of food and light is measured in units of carbon as *ρ*_C:F_ = *ρ*_C:L_ = 1).

Note that we characterise a cells resource uptake ability by the affinity *a_X_*, following Aksnes and Cao (2011); Fiksen et al. (2013); Flynn et al. (2018). This choice contrasts the commonly used Monod/Michaelis-Menten formulation of the functional response, where uptake is described with a half-saturation coefficient and a maximum uptake rate. In a mechanistic context, the Monod formulation of the uptake rate is problematic because the half-saturation coefficient cannot be associated with a physical or physiological characteristic of the cell – it acts purely as a convenient fitting parameter. Mathematically, the affinity follows from the Monod formulation as the product of the half-saturation coefficient and the maximum synthesis rate, which we use to relation to calculate affinities from literature sources of half saturation coefficients. The Monod formulation also includes the process of saturation, which we return to later. Separating the processes of encounter and biosynthesis explicitly with two different parameters (affinity and maximum synthesis rate) avoids the pitfalls of considering the half-saturation constant as a physiological trait (Kiørboe and Andersen, 2019).

The affinity *a_X_* measures the cell’s ability to encounter and assimilate resource *X*. The affinity is determined partly by encounter with the resource and partly by the cell’s investment in capacity to take up and assimilate the resource (Shuter, 1979; Bruggeman and Kooijman, 2007; Chakraborty et al., 2017). The encounter results from the physical processes of diffusion, self-shading, and fluid dynamics. The limitation due to uptake capacity is relevant when the cell encounters abundant amounts of the resource but is unable to process it all by its uptake machinery, e.g., porters for diffusive uptake, light harvesting machinery, or phagotrophic assimilation. For all resource uptakes, the mass-specific affinity is constant or decreases with size, as we will show below. Uptake limitation is most prominent for small cells that have high affinities, leading to a higher encounter with resources that they can process. If investments in uptake capacity scales with the mass of the cell, the uptake limitation of mass-specific affinity is independent of size. Small cells therefore have limited ability to increase their uptake capacity, and their affinity will be limited by uptake capacity. The affinity therefore has two size-scaling regimes: for small sizes the affinity is independent with size (uptake limitation), and for larger sizes it is constant or declining with size (encounter limitation) (e.g. Armstrong, 2008).

The processes that determine encounter and uptake capacity, and how they scale with cell size, depend on the type of resource.

#### 3.1.1 Encounter and uptake of dissolved matter

The theory behind uptake of dissolved matter is well developed, as reviewed by Fiksen et al. (2013). Nutrient uptake is limited by three processes: the rate at which molecules diffuse towards the cell, the rate at which nutrients are transported across the cell membrane by porters, and the capacity of the cell to utilize nutrients in biosynthesis.

The flux of molecules towards a sphere was shown by Pasciak and Gavis (1974) to be proportional to the sphere’s radius and the difference between the concentration far away and at the surface of the sphere. Assuming that the sphere absorbs all encountered molecules the concentration at the surface is zero and the mass-specific affinity becomes:

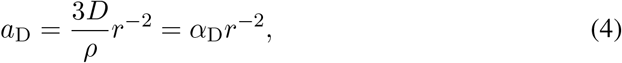

where *D* is the diffusivity of the dissolved molecules, *ρ* is the cell carbon density, and *r* is the cell radius. However, if the cell is embedded in an external flow, such as turbulence or when the cell is sinking, then the boundary layer around the cell will be smaller and the flux of molecules increased. The increase in flux due to such advective flows is characterized by the Sherwood number, which is the dimensionless ratio between transport by advection and diffusion (Kiørboe, 1993). A Sherwood number *≫* 1 means that the transport is enhanced by advection. However, Kiørboe (1993) found that for most cases the Sherwood number is very close to 1, such that Eq. 4 does not have to be corrected for advective effects.

The simple scaling in of affinity in Eq. 4 has formed the start of an extensive theoretical discussion of additional effects cell size on the affinity (reviewed by Fiksen and Jørgensen, 2011). We provide a full mathematical derivation in Box 1 and proceed with qualitative arguments here. At small cell radius, where the mass specific affinity is very high, uptake might become limited by either the number and capacity of porters, or by the cells’ ability to process incoming nutrients. Berg and Purcell (1977) accounted for uptake limitation by introducing an extra term in Eq. 4 (see Box 1 for complete derivation and discussion):

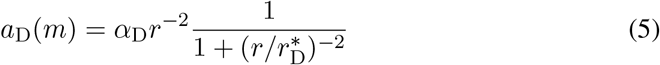

where 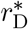 is the cell size at the cross-over between uptake (porter/processing) limitation and diffusion limitation. Small cells 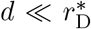 are porter/processing limited with affinity 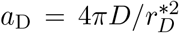 while larger cells, 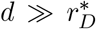 are diffusion limited with *a*_D_ = 4*πDr^−^*^2^ (Fig. 2). Precisely what controls the cross-over size 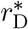 remains uncertain. Geometric consideration based on the size and density of porter site on the cell have been explored (Casey and Follows, 2020; Armstrong, 2008) as have the kinetics of porter handling times and energy costs (Aksnes and Egge, 1991) (Box 1) but remain unresolved.

**Figure 2:**
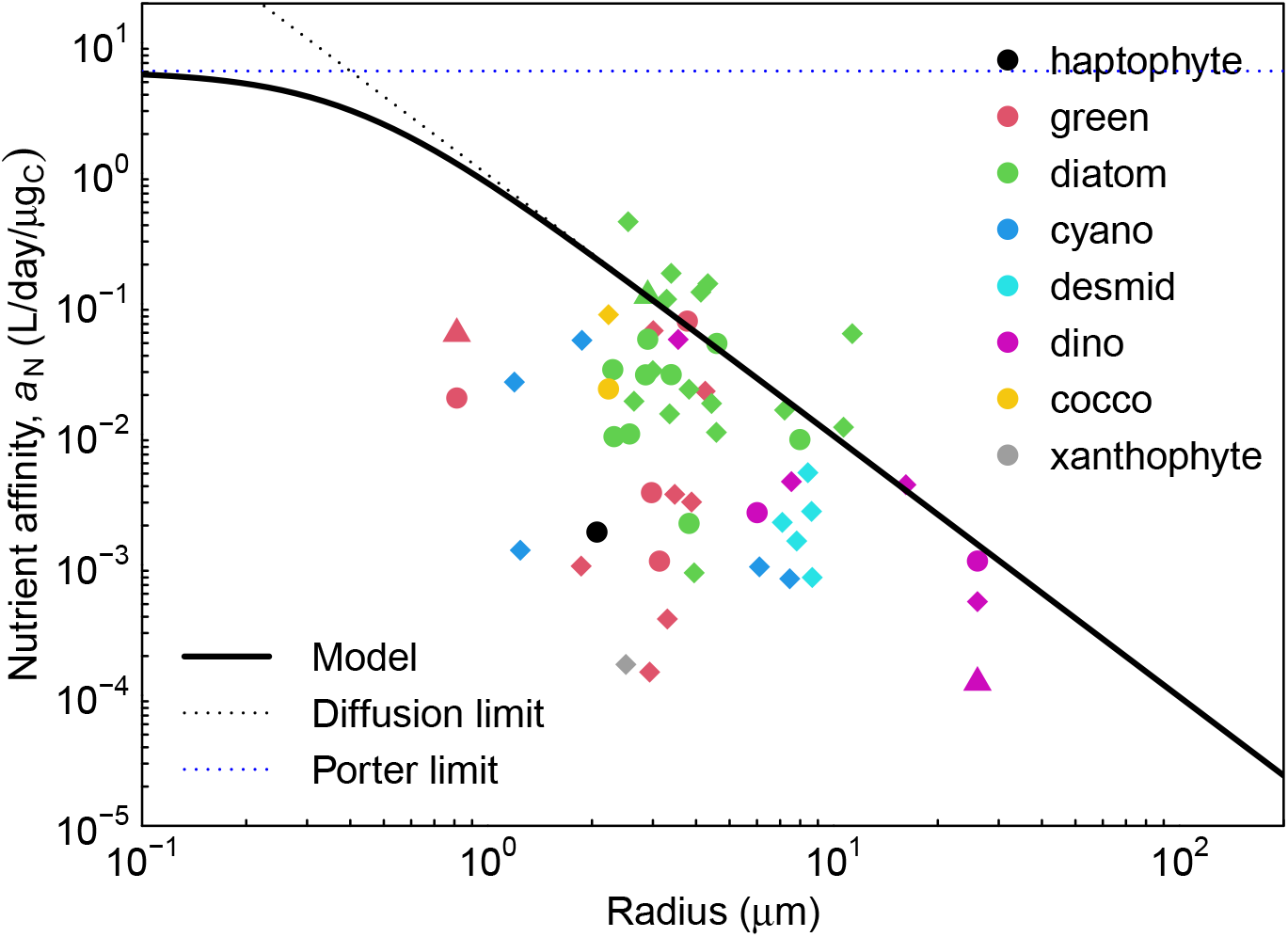
Specific nutrient affinity (*a*_N_) as a function of radius. Triangles: ammonium uptakes; circles: nitrate uptake; diamonds: phosphorous uptake. The dotted lines are the theoretical maximum affinity due to diffusion limitation and porter limitation. The solid line is a fit-by-eye of the radius where porter limitation becomes important, around 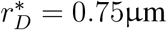. Data from Edwards et al. (2015). Conversions between volume and mass are done using the relations in Menden-Deuer and Lessard (2000).

As a practical solution to determine the cross-over size between diffusive encounter limitation and uptake limitation, we turn to observations. The available data are, however, very scattered (Fig. 2; see Table 1 for a summary of all parameters). The data do confirm the theoretical prediction of an upper limit to encounter by diffusion limitation. The data also indicate that the affinity of smaller cells (smaller than around 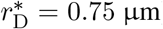) is limited by another process than diffusion limitation, which could be porter or uptake limitation. It should be noted, though, that the date do not lend much support to the value of *r*_D_, only that it should not be less than 1 μm. Below these two upper limits there is a large scatter in the data with some species having a factor 1000 smaller affinity for phosphorous. Our interpretation of this scatter is that species adapted to high nutrient loads, like the fresh-water green algae, are not diffusion or porter limited. They therefore invest less in nutrient uptake with the result that the affinity is smaller than it could potentially be. In the following we use the diffusion/porter limitation to define nutrient affinity as it well represents the affinity in communities with strong nutrient competition. In other communities, e.g. during a spring bloom where nutrients are plentiful, it does not matter that this formalism predicts a too affinity as growth will be limited by the ability to perform biosynthesis and not by nutrient uptake. In conclusion, we have a fully developed theoretical apparatus to understand the maximum affinity of cells to dissolved organic matter, however, we need a better understanding of the specific processes related to molecule capture to fully relate the limitation at small cell sizes to fundamental processes.

**Table 1:** Parameters for the cell-level processes (Section 3) and the size-based model (Section 4.5). “d” is the unit of time (days).

| Parameter | Value | Reference |
| --- | --- | --- |
| *Cell size and density* |  |  |
| Carbon density | $\rho = 0.4 \cdot 10^{-6} \mu\text{g}_C/\mu\text{m}^3$ | (a) |
| Membrane thickness | $\delta = 50 \text{ nm}$ | (f) |
| C:N mass ratio | $\rho_{C:N} = 5.68 \text{ g}_C/\text{g}_N$ | |
| *Cell rate parameters* |  |  |
| Diffusive aff. coef. | $\alpha_D = 0.972 \text{ l} \cdot \mu\text{m}^2/\text{d}/\mu\text{g}_C$ | Fig. 2, Box 1 |
| Diffusive aff. cross-over | $r_D^* = 0.4 \mu\text{m}$ | Fig. 2, Box 1 |
| Light aff. coef. | $\alpha_L = 0.3 (\text{d} \cdot \mu\text{mol m}^{-2} \text{s}^{-1})^{-1} \mu\text{m}$ | Fig. 3, Box 2 |
| Light aff. cross-over | $r_L^* = 7.5 \mu\text{m}$ | Fig. 3, Box 2 |
| Light uptake eff. | $\epsilon_L = 0.8$ | |
| Clearance rate | $a_F = 0.018 \text{ l}/\text{d}/\mu\text{g}_C$ | Fig. 4, Box 3 |
| Max. phagotrophy coef. | $c_F = 30 \mu\text{m}/\text{d}$ | Fig. 5 |
| Assimilation efficiency | $\epsilon_F = 0.8$ | |
| Passive loss coef. | $c_{\text{passive}} = 0.03$ | Sec. 3.2 |
| Max. synthesis coef. | $\alpha_{\text{max}} = 1.5 \text{ d}^{-1}$ | Fig. 5 |
| Basal metabolism coef. | $\alpha_R = 0.1$ | |
| *Prey encounter* |  |  |
| Predator-prey mass ratio | $\beta = 500$ | (b) |
| Predator-prey width | $\sigma = 1.3$ | (b) |
| *Community model parameters* |  |  |
| DOC remin. of feeding | $\gamma_F = 0.1$ | (e) |
| DOC remin. of lysis | $\gamma_v = 0.5$ | (e) |
| Lysis mortality coef. | $\mu_{v0} = 0.004/\log(m^+/m^-)$ | (c) |
| Size of HTL mort. | $m_{\text{htl}} = 8.9 \cdot 10^{-5} \mu\text{g}_C$ | (d) |
| HTL mortality coef. | $\mu_{\text{htl}0} = 0.1 \text{ d}^{-1}$ | Sec. 5.2 |
| *Chemostat parameters* |  |  |
| Mixing rate | $d = 0.0001 \dots 1 \text{ d}^{-1}$ | Sec. 4.5 |
| Deep nutrient conc. | $N_0 = 50 \mu\text{g}_N/\text{l}$ | Sec. 4.5 |
| Productive layer | $M = 20 \text{ m}$ | |

**Box 1: derivation of nutrient affinity**

The diffusive flux of a substance *N* to a partial absorbing sphere of radius *r* has a well known solution:

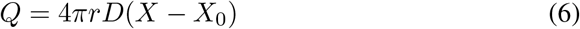

where *D* molecular diffusivity, and *N* and *N*_0_ the nutrient concentration at distance and on the cell surface respectively. For a perfectly absorbing sphere *N*_0_ = 0 and the flux becomes *Q* = 4*πrDN*. A real cell however is not perfectly absorbing but is covered by a finite number of uptake sites in an otherwise impervious cell membrane. A classic result (Berg and Purcell, 1977) considers the cell surface is covered by *n* porter sites each of radius *s*. If sites are small (specifically *s ≫ r*), sparsely distributed, and perfectly absorbing, then the diffusive flux towards each site is *Q_s_* = 4*sDN*_0_. For *n* such sites then

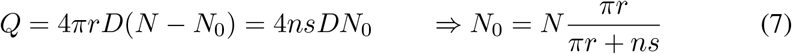

which leads to:

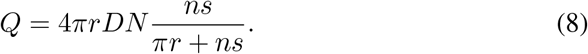

A correction accounts for potential interference of diffusive fluxes when porter sites are tightly packed (Zwanzig, 1990). Specifically, expressing the surface fraction of porters as *p* = *ns*^2^/(4*r*^2^)

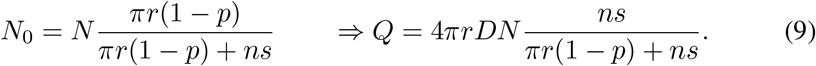

As *p →* 1 (i.e. the entire cell surface becomes covered with perfectly absorbing porter sites) *N*_0_ *→* 0 and *Q →* 4*πrDN*. While theoretically sound and widely built upon, these results are actually not particularly germane to the question of nutrient uptake in plankton. In the first instance, for typical cell sizes and porter sizes, the correction (Eq. 9) saturates extremely rapidly so a very low porter density is sufficient to achieve near maximum uptake flux (Jumars et al., 1993). This implies that limitation of the number of porter sites due to surface crowding is unlikely to be an issue. Secondly, it is not realistic that uptake sites are perfectly absorbing discs. While diffusion towards the sites is a fair representation, uptake requires active transport across the cell wall (Aksnes and Egge, 1991; Armstrong, 2008), a process that (1) occupies the uptake site for a finite amount of time and (2) is energetically costly, requiring about 1 mole of ATP per mole of nutrient transported.

#### 3.1.2 Light harvesting; theory and data

**Box 2: derivation of light affinity**

The net absorption of light by a cell depends on the density and distribution of individual chromophores within the cell’s cytoplasm. For a spherical cell (radius *r*) with uniformly distributed chromophores throughout the cell volume (number density *c* (μm^−3^), optical cross section *a* (μm^2^)), the rate at which photons are absorbed is given by:

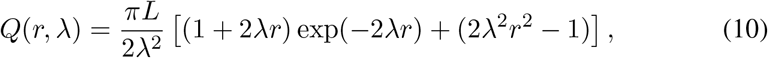

where *λ* = *ac* (μm^−1^) is the light absorption coefficient within the cytoplasm, and *L* (μmol m^−2^s^−1^) is the light flux (Duyens, 1956; Kirk, 1975). This relationship, while exact for a sphere, is somewhat clumsy. A more accessible formulation, developed by Hansen and Visser (2019), assumes a cylindrical cell with the same volume and cross-sectional area as the sphere. Under this geometry, the optical path through the cell is 4*r*/3 and:

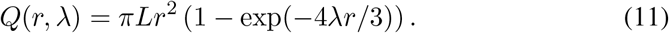

Given that these formulae give very nearly identical results, and that cell shape is always a confounding factor, we opt for the simpler. From the form of (Eq. 11) it is clear that for large cells with a high investment in chromophores (*λr ≫* 1), photon absorption is proportional to the cell’s cross-sectional area *Q ≈ πr*^2^*L* whereas for small cells with low chromophores investment (*λr ≫* 1), photon absorption is proportional to cell volume *Q* ≈ 4/3*πr*^3^*λL*.

The mechanisms relating photon absorption to carbon fixation are complex and dependent a variety factors including photon energy, type of pigments and details of the photosystem used. While some of these aspects are accessible to modelling, we use the commonly used quantum yield *y* (gC/(mol photon)) as a simplification (Emerson, 1958). The specific light affinity then becomes *a*_L_ = *y*(*Q/L*)/*m*. We can also write *λ* = *κ*_L_*φ*_L_ relating the cell’s absorption coefficient to *φ*_L_, the fraction of its carbon mass invested in light harvesting where *κ*_L_ the constant of proportionality. Observations indicate that *λ* = 0.1 μm^−1^(Raven, 1984, 1997) when about half of the cell’s mass is devoted to light harvesting, suggesting that *κ*_L_ = 0.2 μm^−1^. It follows then that

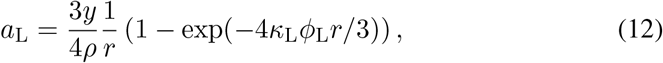

which is identical to Eq. 13 with parameters 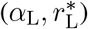 corresponding to (3*y*/(4*ρ*), 3/(4*κ*_L_*φ*_L_)) respectively. Quantum yield estimates ranges from 0.12 to 0.6 g_C_/(mol photon) (Kishino et al., 1986). Using *y* = 0.16 gC/(mol photon) suggests *α*_L_ = 0.30 (d μmol m^−2^s^−1^)^−1^μm and 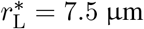.

Fig. 3 shows that there are cells with almost a factor 10 higher affinity that predicted by Eq. 12. The source of this variation is likely due to uncertainty in the quantum yield *y*, which depends on the type of pigment and the wavelength of the light (Kishino et al., 1986).

**Figure 3:**
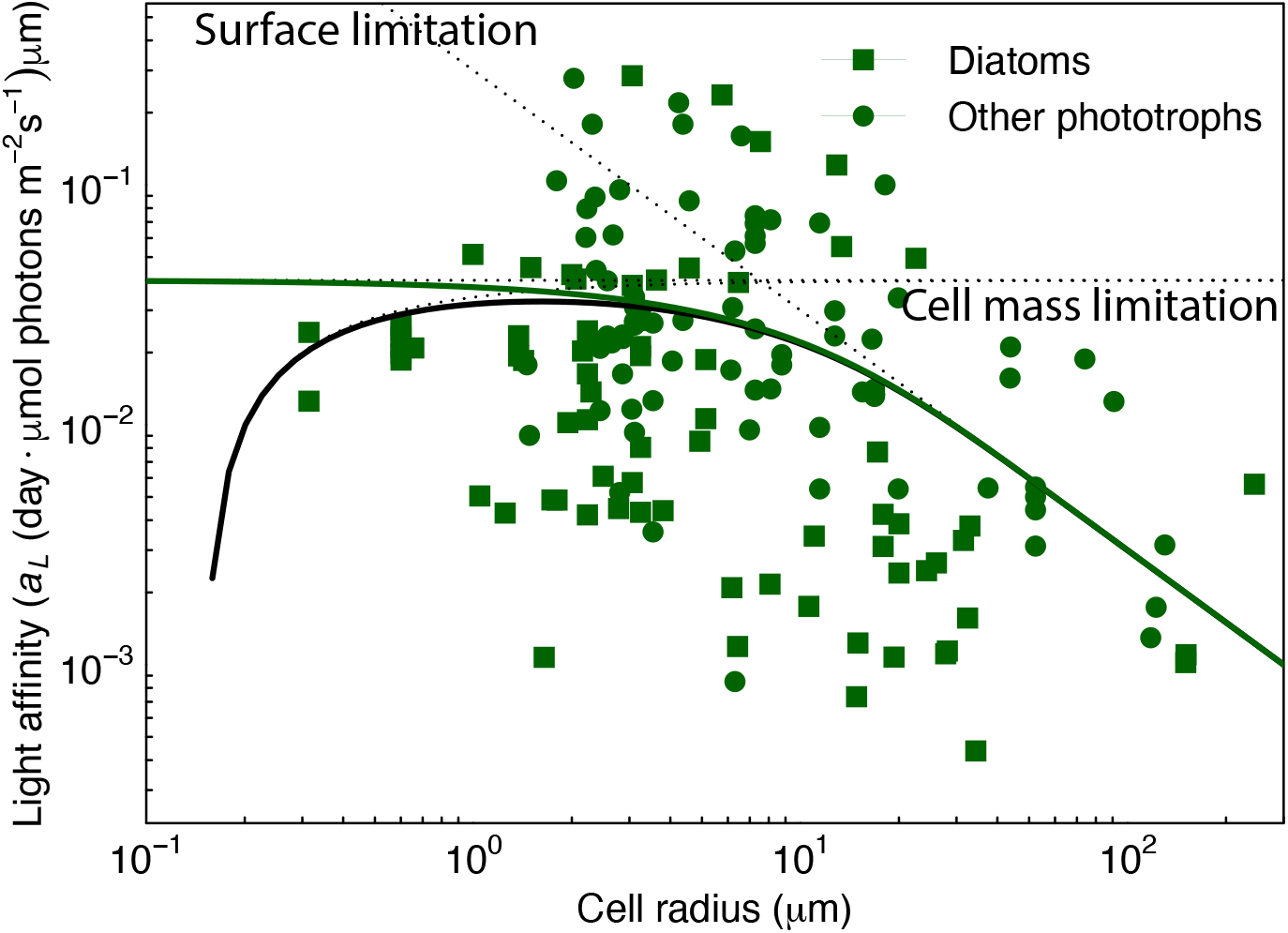
Light affinities of protists as a function of carbon mass compared to the first-principles formulae (Eq. 13; thick black line and (Eq. 12); green line). The three limiting factors: cells mass, cell surface, and cell membrane are shown with dotted lines. The black line is the total affinity. Data from Edwards et al. (2015), corrected for day length.

Photosynthesis is fundamentally powered by the capture of photons by light harvesting complexes, and the number of photons captured by a cell depends on both the number of photons incident on the cell, as well as the number of light harvesting complexes within the cell. In terms of scaling, it can be reasoned then that the former depends on the cross sectional area of the cell, while the latter on some proportion of its functional carbon mass. However, light harvesting complexes shade one another and in larger cells not all complexes can be equally effective for light harvesting (Kirk, 1975; Morel and Bricaud, 1981). The affinity for light harvesting therefore transitions from being independent of size for small cells to being proportional to the surface area for large cells (see Box 2):

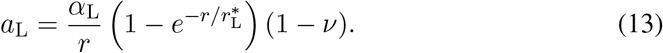

This formulation of affinity has asymptotic scaling of 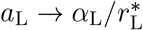 for intermediate cells, *a*_L_ *→ α*_L_/*r* for 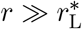 and goes to zero for small cells (the factor 1 *− ν*).

Previous analyses of light affinity has focused on fitting just one power law and has consistently found a scaling close to the predicted surface law *∝ r^−^*^1^ (Taguchi, 1976; Finkel et al., 2010; Edwards et al., 2015). Our reanalysis of the available data indicates a transition from mass to surface scaling with a transition size around 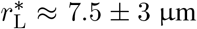 (Fig. 3) in accordance with the first-principles argument in Box 2. The divergence to zero due to the cell wall limitation 1 *− ν* for smaller cells is consistent with a lower limit of one chloroplast at cell volume of ≈ 1 μm^3^ or *r* ≈ 0.5 μm(Okie et al., 2016).

As with the nutrient affinity there is a large scatter in the data of one order of magnitude around the first-principle prediction. Here, though, the prediction does not reflect the upper limit of light affinity rather an average estimate. This indicates that some plankton can invest more in light harvesting to increase their affinity. In developing the prediction we assumed that plankton invest at most half of their cell mass to light harvesting. Cells might invest more if they are fully dedicated to light harvesting in low light environments leading to a higher affinity. Further, the quantum yield is uncertain and a higher value is within the observed range. However, the absolute value of the affinity is less important for the plankton community than how affinity scales with cell size. In a water column, the production maximum adjusts itself vertically to the point where light limitation matches nutrient limitation (Ryabov et al., 2010). Therefore, a higher light affinity leads to a deeper production maximum and vice versa. The overall value of the affinity is therefore less important for the general production of the plankton community because production will be limited by nutrients, unless the light is so low that production can only occur in the surface. What is important, though, for the structure of the trophic strategies with cell size is that the specific affinity decreases with cell size overall, and that decline is well borne out by the data.

#### 3.1.3 Phagotrophy

**Box 3: Derivation of clearance rate**

The specific clearance rate *a*_F_ (volume/time/cell mass) can be estimated from the work required to displace the fluid that a cell moves through or filters. We assume that the work is approximately the same as pushing a sphere through the fluid, i.e., given by Stokes’ law: *W* = 12*πμu*^2^*r*, where *u* is the velocity, *r* the cell radius, and *μ* the dynamic viscosity of water. The metabolic power that the cell has available to filter water scales with the cell’s mass *cmρ*_e_, where *c* is the fraction of the cell’s mass that can be used for swimming and *ρ*_e_ is the energy density of the cell. Equating the work needed and the power available gives the velocity as:

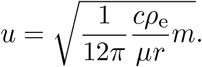

Assuming that the cell clears an area corresponding to its own cross section we get the specific clearance rate as the clearance area *πr*^2^ multiplied by the velocity and divided by the mass:

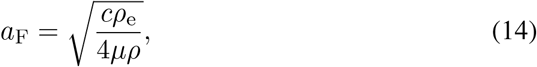

using Eq. 1 to convert between radius and mass. The specific clearance rate is constant (independent of cell size). With a dynamic viscosity of *μ* = 1 g/(m s), energy density *ρ*_e_ = 40 *·* 10^3^m^2^*g*s^−2^g_C_^−1^ (Boudreau and Dickie, 1992), and that the fraction of body mass used for driving the flow is 0.1 day^−1^ gives *a*_F_ = 0.0073 l/day/μg_C_, very close to the geometric average of 0.018 from the data in Fig. 4.

**Figure 4:**
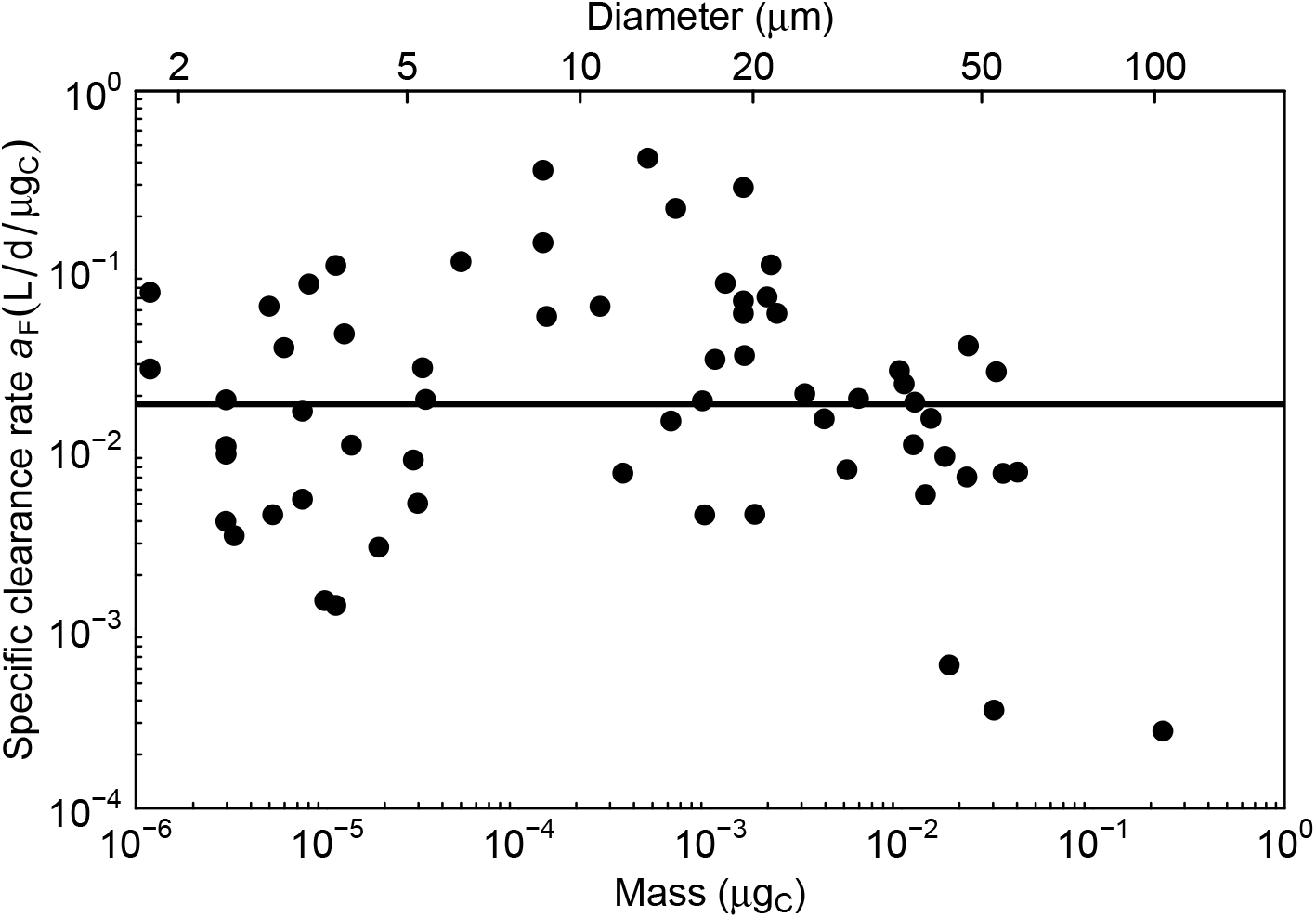
Specific clearance rate (*a*_F_) as a function of carbon mass. Data of nanoflagellates, dinoflagellates, and ciliates from Kiørboe and Hirst (2014).

Phagotrophy is the ingestion of food particles, typically smaller cells. Prey cells are encountered either by the predator moving through the fluid or with the predator creating a feeding current that brings prey towards it (Kiørboe, 2011). In this case the affinity is the clearance rate, i.e., the volume of fluid cleared of potential prey per time. Hansen et al. (1997) showed that the half saturation constant and the maximum consumption rate was roughly constant among unicellular plankton, which corresponds to a constant mass specific clearance rate. Kiørboe (2011) expanded the analysis a showed that the clearance rate was approximately 10^6^ cell volumes per day, though variations exist among feeding modes (passive, active, cruising or feeding current). It appears evident that the scaling of clearance rate with cell size should emerge from fluid mechanic constraints. Despite arguments having been made for fish, they have not been made for unicellular plankton. We develop an argument in Box 3 that reproduces the observed constant specific clearance rates and also gets the average value reasonably correct (Fig. 4).

For the actual food consumption, we also need to consider the limitation imposed by assimilation over the food vacuole membrane. The surface area of the vacuole scales *∝ r*^2^ and the specific maximum assimilation therefore scales with *r^−^*^1^. We can then described the uptake with a classic functional response with affinity *a*_F_ and maximum assimilation rate *c*_F_*r^−^*^1^:

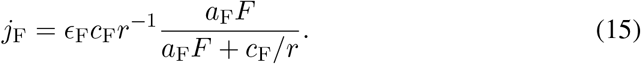

where *E*_F_ is the assimilation efficiency. This formulation has the limit *j*_F_ *→ E*_F_*a*_F_*F* for smaller cells and *j*_F_ *→ E*_F_*c*_F_/*r* for larger cells with the cross-over size between the two regimes being food-dependent: 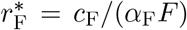. We do not have any direct measurements of the assimilation limitation, however, we will use the measurements of maximum growth rate of larger cells to estimate this process as *c*_F_ ≈ 30 μm/day. It can be argued that the reduction in functional mass of small cells (the factor *ν*) should lead to a reduction in phagotrophy for small cell, similar to the reduction in phototrophy. However, phagotrophy is not relevant for the smallest cells because they have no suitable food, so including the effect of cell membrane for phagotrophy is irrelevant.

### 3.2 Passive losses across the membrane

It is well recognized that cells leak smaller molecules across their membrane, however, the exact processes behind this loss are not well understood. Bjørnsen (1988) distinguished between losses as “income taxes” and “property taxes”. Income taxes are those losses incurred during uptake. These losses are represented as a less than 100% efficiency of the uptakes. Property taxes are those losses that occur regards of the uptakes, which we here consider as passive exudation. The passive exudation can be assumed to scale with the surface area (Kiørboe, 2013) and, assuming a negligible external concentration, becomes:

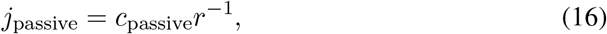

where *c*_passive_ = 3*P* where *P* is the permeability of a phytoplankton membrane. Values of the membrane permeability varies wildly: Braakman et al. (2017) argues for a very high membrane permeability in excess of ≈ 10^6^ μm/day. This high permeability would imply that the cell spends significant amounts of energy continuously re-uptaking lost nutrients. Bjørnsen (1988) considers that *P* ≈ 1 μm/day. Even this value is very high. However, considering than only about 10% of the compounds are of sufficiently low molecular weight to escape through the membrane, *c*_passive_ should be reduced by a factor 10. Further, the smallest cells, which are those which are most affected by passive exudation losses, are bacteria with a different cell membrane than the phytoplankton considered by Bjørnsen (1988). We therefore propose to further reduce the permeability further and use *c*_passive_ ≈0.03 μm/day.

### 3.3 Biosynthesis and basal metabolism

The maximum rate of biosynthesis is limited by the cell’s investment in synthesis machinery, i.e., ribosomes. If we consider the number of ribosomes to be proportional to the functional cell mass then the synthesis rate, the biomass synthesized per time and per cell mass, becomes independent of functional cell mass, i.e., *∝* 1 *− ν*:

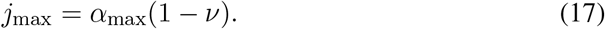

We have no first-principle arguments to set the level of maximum synthesis rate *α*_max_ (but it may be possible to develop an argument based on the size and capacity of a ribosome). A more detailed argument that dynamically predicts the maximum synthesis rate as a trade-off between investment in ribosomes and chloroplast has been developed (Shuter, 1979; Serra-Pompei et al., 2019, e.g.), however, even then the crucial parameters are not constrained by first principles arguments constrained. The available data show a large scatter with maximum synthesis rates varying between almost zero and 3 day^−1^ (Fig. 5). The data also indicates that maximum synthesis rates are lower for small and large cells than for intermediate-sized cells. The reduction in max synthesis rate of large cells can be explained by the limitation due to phagotrophic assimilation (Eq. 15) as larger cells are purely phagotrophic. We have, rather arbitrarily, chosen a value of *α*_max_ = 1.5 day^−1^. This value does not represent the upper limit and it will therefore somewhat limit the community’s ability to create a strong bloom in a seasonal environment.

**Figure 5:**
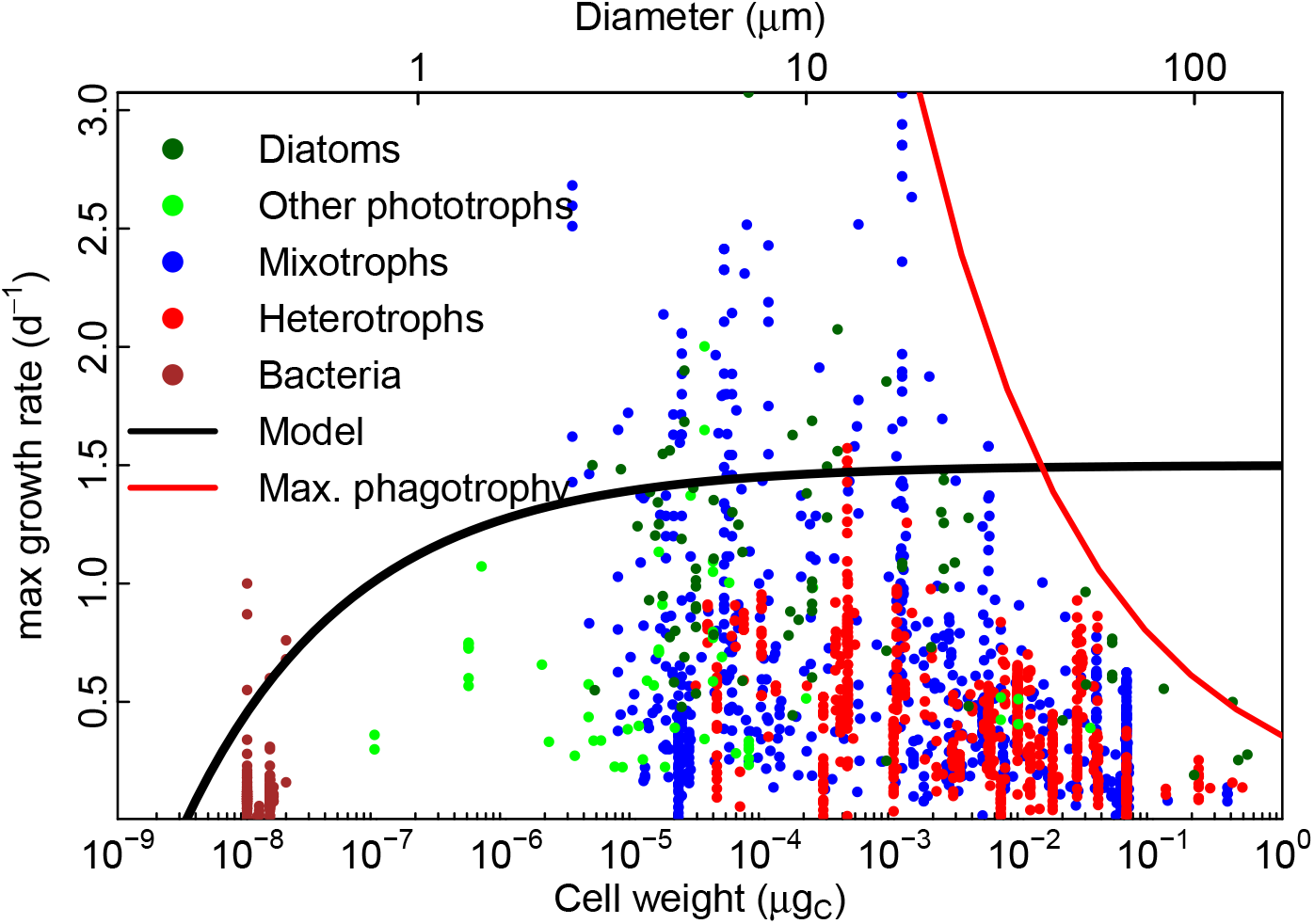
Maximum growth rates of plankton. Phototrophs from Edwards et al. (2015), mixotrophs (nano- and dinoflagellates and ciliates from Kiørboe and Hirst (2014) and “bacterivores” from Rose and Caron (2007)), heterotrophs (“herbivores” from Rose and Caron (2007)) and of bacteria (Kirchman, 2010). Rates are converted to 10 degrees with a *Q*_10_ = 1.5 for phototrophs and *Q*_10_ = 2.8 for mixo- and heterotrophs. The solid line is Eq. 17 with *α*_max_ = 1.5 day ^−1^. The red line is the maximum assimilated phagotrophic uptake *E*_F_*c*_F_/*r*. The diameter-axis on the top of the panel is not accurate for diatoms because of their vacuole which gives them a smaller density than other cells.

The division rate is further limited by the basal metabolism. The basal metabolism supports the functions needed to keep the cell alive but not the respiration associated with resource assimilation and biosynthesis. In this simple model we do not distinguish between basal metabolism and other respiration (but see Chakraborty et al. (2017, 2020)) and consider simply that all respiratory costs are a fraction of cell mass, and therefore that *j*_R_ is constant. For simplicity we write it proportional to the maximum synthesis capacity:

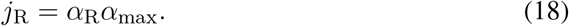

with *α*_R_ ≈ 0.1.

### 3.4 Temperature effects

The temperature response of a cell is commonly modelled by multiplying the maximum growth rate with a *Q*_10_ or Arrhenius factor. For heterotrophic plankton a *Q*_10_ ≈ 2 well represents the temperature response of cell metabolism, whereas a lower factor is used for phytoplankton. It is therefore common for models to use different *Q*_10_ factors for phyto- and zooplankton (e.g. Archibald et al., 2022). However, the temperature response of phototrophic plankton is more complex, and recent experimental work has shown a strong dependence on the resource environment (Schaum et al., 2017; Thomas et al., 2017; Marañón et al., 2018). Shuter (1979) showed how temperature effects in phytoplankton should emerge as a result of the *Q*_10_’s of each metabolic or resource uptake process in the cell. Serra-Pompei et al. (2019) took this idea further and applied it to mixotrophic plankton. They found that temperature responses of the cell’s growth rate varied between almost no temperature response in environments with low nutrients and high light, to around *Q*_10_ = 2 in high food environments.

Temperature effects are introduced by multiplying rates with a *Q*_10_ function: 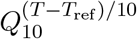 which gives the fractional increase in the rate when the temperature is increased 10 degrees from a reference temperature of *T*_ref_ = 10 degrees. The *Q*_10_ factors are (Serra-Pompei et al., 2019): *Q*_10_ = 1.5 for diffusive uptakes, no temperature correction for light capture and a standard “metabolic” correction of *Q*_10_ = 2 on respiration, maximum phagotrophy, and maximum synthesis capacity. For feeding one could follow Eq. 14 and use the temperature scaling of viscosity (*Q*_10_ ≈ 1.5). However, prey also have escape maneuvers which will become equally faster so we assume that the two effect cancel one another and use *Q*_10_ = 1 for feeding.

## 4 Size structure of the plankton community

The previous section was devoted to describe the processes of the single cell as a function of its size and tie this processes down to first principles as far as possible. This section is devoted to analyse the structure of the plankton community and how it emerges from the first principles constraints on the cell processes. What actually defines the “size structure” of a community? It is how the community varies with cell size: which types of cell dominate a given size group and how big is their biomass.

The section is split into two parts: first we analyse the cell’s resource uptake and metabolism as a function of size to identify the maximum and minimum size of cells, the competitive abilities of different sized cells, and their dominant trophic strategies. In the second part we scale from the cell-level processes up to the biomass distribution of the plankton community, both with a simple theoretic argument and with a full dynamic model.

### 4.1 Smallest and largest cells

Raven (1994) argued that the cell membrane sets a lower limit of the size of the smallest cell. The absolute smallest size is when the cell membrane uses the entire mass, i.e., when the cell membrane fraction *ν* = 1 (Eq. 2):

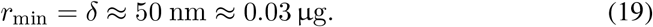

This is an extreme lower limit for a cell with plenty of resources and no losses. Considering that losses to respiration and passive losses (Eq. 16) can not exceed the maximum synthesis rate (Eq. 17): *j*_max_ > *j_R_* + *j*_passive_, gives a larger minimum size of:

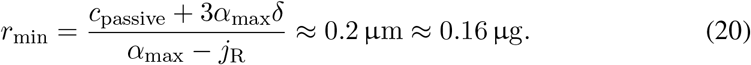

The largest unicellular plankton are heterotrophs (Andersen et al., 2016). They are limited by two processes: the rate at which oxygen diffuses into the cell (Fenchel, 1987; Payne et al., 2011) and the rate at which they can assimilate food through their feeding vacuoles (red line in Fig. 5). Considering the limiting effects of food uptake, the maximum size *r*_max_ is when the maximum rate of assimilated consumption, *E*_F_*c_F_ /r* (Eq. 15) equals the metabolic costs *j*_R_:

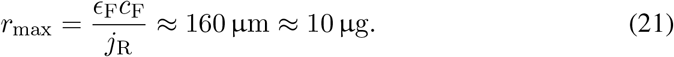

Fenchel (Chap. 1 1987) considered that the upper size limit is imposed by the diffusion of oxygen into the heterotrophic cell. He finds that the largest radius where O_2_ diffusion can satisfy the metabolic demand is:

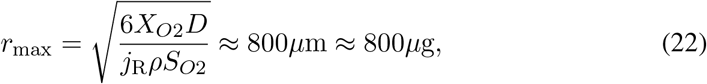

where *X_O_*_2_ is the external oxygen concentration, *D* the diffusivity of oxygen, and *S_O_*_2_ the oxygen:carbon mass ratio (Payne et al. (2011) did a similar evaluation and found an upper limit around 1 mm under present day oxygen concentrations). The upper limit imposed by oxygen is rather large compared to the upper limit imposed by assimilation (Eq. 21) and it is tempting to disregard oxygen as a constraint on maximum cell size. However, it is instructive to look also on the size distribution on cell shape, as analysed by Ryabov et al. (2021) (Fig. 6). The smallest cells are spherical, which is the shape that minimizes the cell membrane per mass. Cells larger than about 0.05 μg_C_are dominated by cylindrical cells. Being cylindrical minimizes the distance of oxygen diffusion from the cell surface to the center. That larger cells are cylindrical therefore indicates the importance of oxygen for the upper limit of cell size. It is possible that not only the diffusion limits the cell size, but also the permeability of the cell wall; a complication that is ignored in the argument by Fenchel (1987).

**Figure 6:**
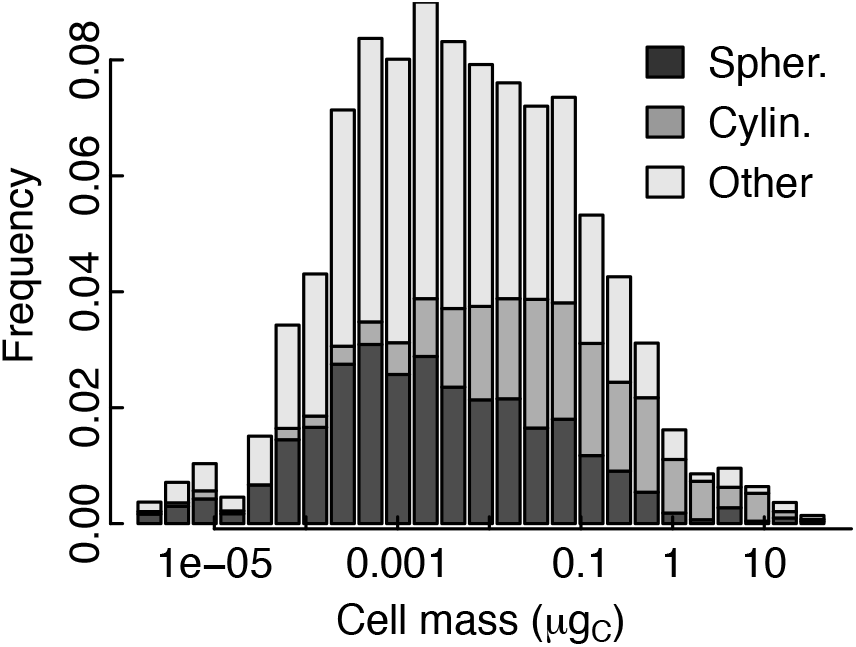
Distribution of cell shapes of phytoplankton as a function of cell mass. Data from Ryabov et al. (2021)

To overcome the upper limitation of size, organisms will have to become multicellular. The smallest adult copepods are on the order of 0.01 μg, which corresponds to the size where the feeding vacuole becomes limiting for growth (Fig. 5).

### 4.2 Limiting resources, R*

The growth of plankton is limited by their ability to acquire and assimilate resources of nutrients, DOC, food from predation, and light. As dissolved resources are subject to competition by all cells, nutrients and DOC are exhausted to the lowest level that the most competitive groups can just survive on. This level is commonly referred to as the “*R^*^*” value, *sensu* Tilman (1982). Ward et al. (2014) calculated the limiting nutrient resource *N ^*^* as a function of cell size and found that limiting resource increases with cell size – confirming the classic result that the smallest cells are the most competitive for nutrients (Munk and Riley, 1952). Here we extend the *R^*^* concept to the concentration of DOC, food, and light. Food is different than the dissolved resources because not all size groups compete for all sizes of food due to size-based selection. Nevertheless, *F ^*^* indicates the minimum level of biomass of their prey. Finally, we can calculate the minimum level of light *L^*^* where purely phototrophic plankton can survive. Plankton does not compete for light (except in extreme cases of biomass as seen in some fresh water environments; Klausmeier and Litchman (2001)), but the *L^*^* indicates the minimum light level – and thus the maximum depth – where photosynthesis alone can support plankton growth.

We can find the limiting resources by calculating the resource level that just balances losses to exudation, respiration, and mortality, e.g., for light: *j*_L_(*L^*^*) = *j*_passive_ + *j_R_* + *μ*, where *μ* is mortality losses (see Table 2). *N ^*^* is calculated from the assumption that carbon is abundant so we can ignore respiration. Similarly, the calculation of *L^*^* and *DOC^*^* assumes abundant nutrients but no alternative carbon source (from DOC, light or food). *F ^*^* also assumes no other carbon source (no phototrophy or DOC).

**Table 2:**
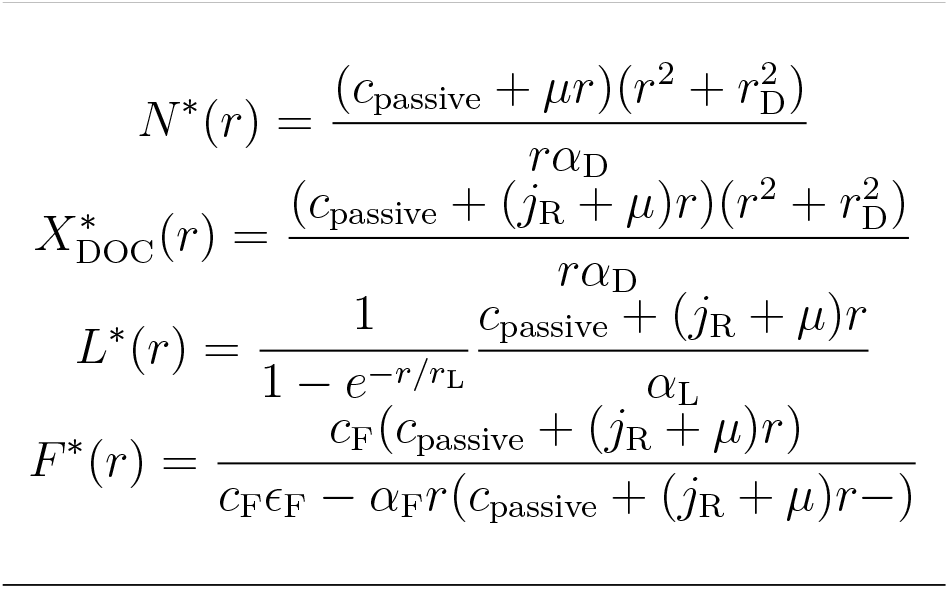
Limiting resource levels

The actual values of the limiting resources cannot be compared directly between one another because they are in different units, however, the interesting aspect is also mainly which size can survive on the lowest resource levels. All limiting resources have a minimum at a specific size (Fig. 7). The minimum emerges as the result of two opposing effects: the passive losses which decreases with cell size (due to decreasing surface to volume ratio; Eq. 16), and the affinity which also decreases (or is constant) with size. The most pronounced minimum is for diffusive uptake of dissolved carbon and nutrients. In contrast to the results by Ward et al. (2014) the *N ^*^* for the very smallest sizes again increases, however, this increase is likely not relevant as the smallest cell are limited by the cell membrane (the grey area in Fig. 7). Regarding light, a very wide range of sizes can survive on the lowest light levels. Phototrophy therefore selects weakly for cell size, and the selection only enters because the cells also need nutrients, which select for small cells. The minimal food requirement *F ^*^* is almost independent of size and is around 1 μg_C_/l. Environments with less food therefore cannot support a longer food chain with purely heterotrophic plankton.

**Figure 7:**
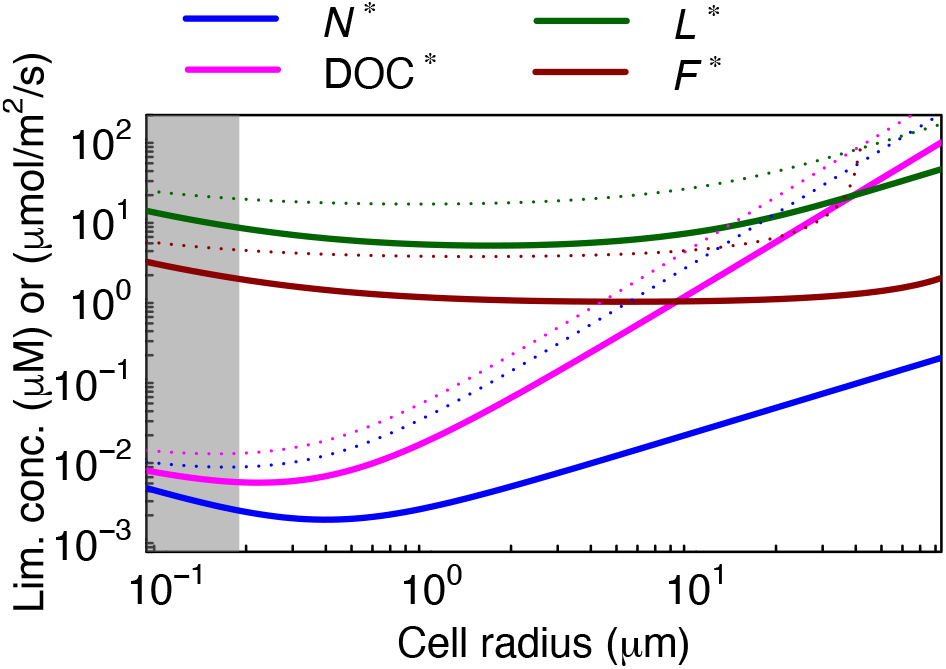
Minimum resource concentrations for survival (*R^*^*). Shown with mortality *μ* = 0 (solid lines) and *μ* = 0.4 day ^−1^ (dotted lines). Carbon sources (light and DOC) assume plenty of nutrients; *N ^*^* assume plenty of carbon; *F ^*^* is for a pure phagotroph. The grey area indicates the minimum viable size of a cell (Eq. 20).

### 4.3 Trophic strategies

The other dimension of community structure is the trophic strategies, i.e., how cells acquire resources: by osmotrophy (diffusive uptake of DOC), phototrophy, or by phagotrophy. The dominant strategy is determined by which of the three fluxes *j*_DOC_, *j*_L_ and *J*_F_ is the largest (Andersen et al., 2016). Fig. 8a shows the fluxes of DOC, carbon from phototrophy, nutrients, and food in an environment specified by concentrations of DOC, N and food (specified by the level of the size spectrum *κ*), and by light. Typically, very small cells are osmotrophs, somewhat larger cells are light-limited phototrophs, medium-sized cells are nutrient limited and larger cells are mixotrophs or heterotrophs. However, the transitions between the dominant strategies occur at different sizes: less nutrients or more available food favours mixotrophic and heterotrophic strategies, while more DOC favours osmotrophy. Andersen et al. (2016) provided analytical expressions for the sizes where the dominant strategies switch from one strategy to the other. This was possible because they used simple power-low relationships for the affinities. Here, however, the relationships are more complex and exact analytical expressions are not possible. However, approximations can be made, which show how the transition sizes depend upon the resource concentrations and the affinities (Table 3).

**Table 3:**
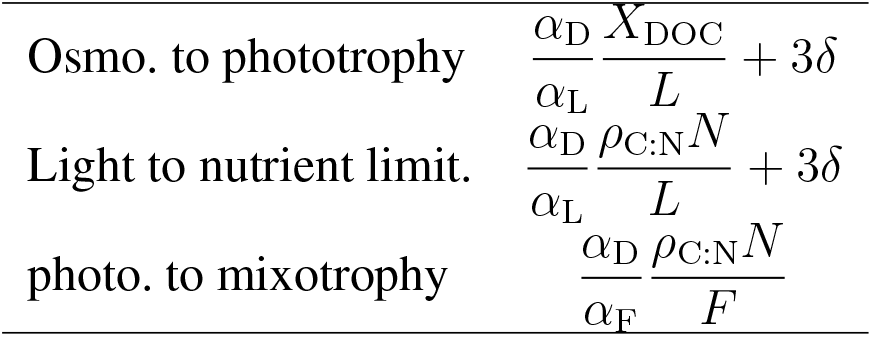
Approximate expressions for the sizes where strategies transition from osmotrophy to phototrophy, from light- to nutrient-limited phototrophy, and from phototrophy to mixotrophy. The expressions are derived by using Eq. 4 for diffusive affinity and by ignoring the correction term in the parentheses of Eq. 13.

**Figure 8:**
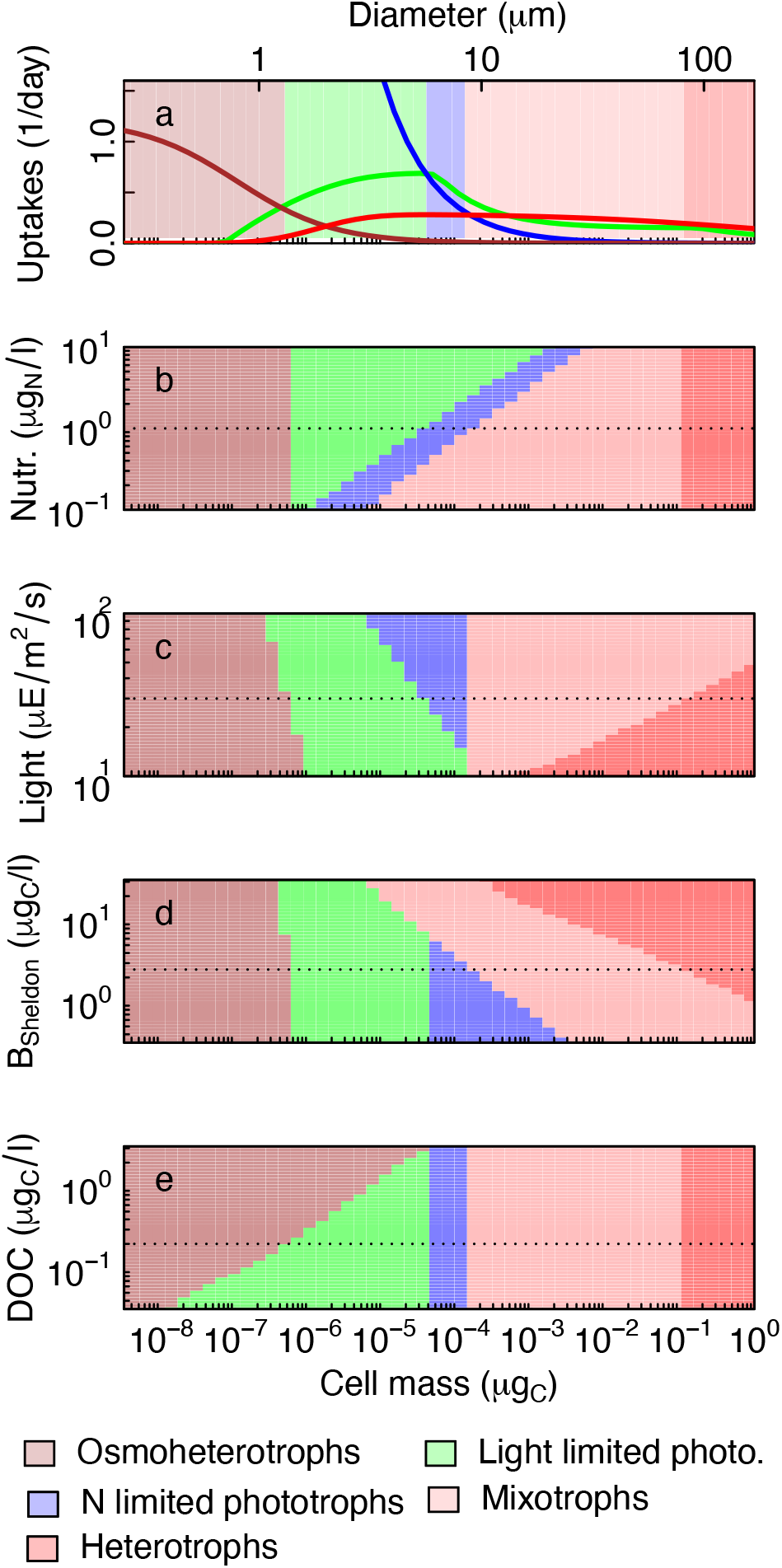
The trophic strategies of unicellular plankton under different environmental conditions. a) The gains from light (green), nutrients (blue), food (red), and DOC (brown), and phagotrophy (red). The dominant trophic strategy is shown by shading: heterotrophy, when surplus nutrient is leaked (red); mixotrophy when the carbon gain from phagotrophy and DOC surpass the potential gain from phototrophy and the nutrient gain from feeding surpass that of diffusive nutrient uptake (light red); nutrient limited phototrophy when the potential gain from phototrophy and DOC surpass the nutrient uptake (blue); light limited phototrophy when nutrient uptake surpass carbon uptake (green); osmotrophy when carbon from DOC surpass carbon from light harvesting (brown). b-e) Variations in the dominant trophic strategy with changes in nutrients, light, biomass and DOC around the conditions in panel (a) are indicated with a dotted line in each panel.

The perspective of trophic strategy being set by the most favourable strategy adds more detail to the argument developed above about the structure being determined by the most competitive size. Generally, the two perspective agree: small cells are dominated by osmotrophs because they are the most competitive for dissolved resources. The perspective of the dominant strategy adds more detail, though, by showing how the smallest phototrophs are light limited while larger phototrophs are nutrient limited, and showing the size ranges of mixotrophs and pure heterotrophs.

**Box 4: Theoretical derivation of the size spectrum**

If the cell is not limited by uptake over the feeding vacuole (i.e., that *a*_F_*F ≫ c*_F_*r* in Eq. 15) then the effective encounter rate is *j*_F_ = *E*_F_*a*_F_*F* (Eq. 15). The encountered food *F* is found by inserting the ansatz *b*(*m*) = *κm^λ−^*^1^ in Eq. 31 to give *F* (*m*) = *κm^λ^α*, where *α* = *√*2*πσβ^λ^e^λ^*2*σ*2/2 is a factor that depends on the parameters of the size preference function (Eq. 30). Following Andersen and Beyer (2006) we now assume that the encounter rate of food *j*_F_ is proportional to the metabolic needs *j*_R_ and independent of size. Then we can equate encountered food with metabolic needs:

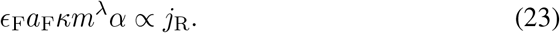

This relation is only true if the dependency on *m* disappears, i.e., if *λ* = 0. When *λ* = 0 the abundance distribution is *b*(*m*) *∝ m^−^*^1^, corresponding to the Sheldon spectrum *B* (Eq. 29) being constant (independent of cell size).

The level of the spectrum, *κ*, can be estimated by assuming that the entire flux of new nutrients *dN*_0_ into the photic zone is taken up and used. This assumption is reasonable as nutrient concentrations in the surface are much less than deep nutrient concentrations in the productive season. The flux of potential new production is *dN*_0_ which can support a new primary production of *dN*_0_*ρ*_C:N_ (g_C_/day/liter). There are three sources of losses: higher trophic level predation, diffusion losses, and respiration. The losses to higher trophic levels are found by integrating over the range where the higher trophic level mortality acts, i.e., a factor *β*:

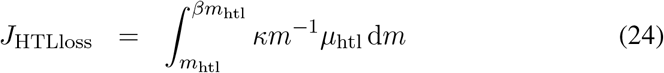

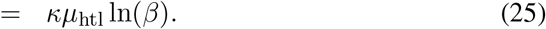

Diffusion and respiration losses are found by integrating over the entire size range:

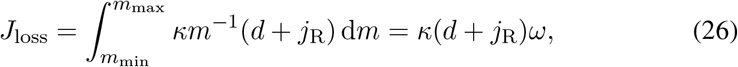

where the ranges of the integration are given by Eqs. 20 and 21, and *ω* = ln(*m*_max_/*m*_min_) ≈ 25. Equating the new production with losses, and accounting for a fraction *E*_htl_ of the higher trophic level losses being remineralized in the photic zone, gives:

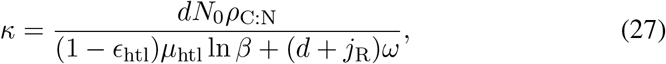

**Box 5: definition of the Sheldon size spectrum**

The structure of the plankton community is represented by the biomasses in the size groups *B_i_*. This representation has the disadvantage that the level of the biomasses depend on the size-range of each group: broader (fewer) size-groups leads to higher average biomass level and *vice versa*. To avoid this dependency size distributions are often shown as “normalized size spectra” (Sprules and Barth, 2016), by dividing the biomass with the size range of the group: 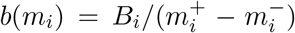, where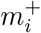 and 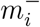 are the upper and lower sizes in the size group. If we assume a scaling biomass spectrum, *b*(*m*) = *κm^λ−^*^1^ then the relation between the normalized biomass spectrum and the binned size groups is:

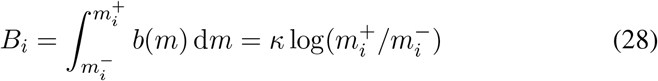

if *λ* = 0. If size groups are evenly distributed on a log scale then 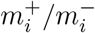 is constant (independent of mass) and the biomasses in each groups are roughly the same. To avoid that results depends on the binning of the size groups we here define the “Sheldon” spectrum as:

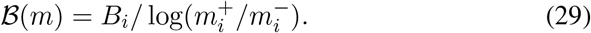

### 4.4 Theoretical size spectrum

The size spectrum was first introduced by Sheldon and Parsons (1967) who plotted the biomass as a histogram in log-spaced size groups and showed that the biomass was roughly independent of cell size. Since that pioneering work the regularity of the log-histogram spectrum has been demonstrated over and over again, as reviewed by Sprules and Barth (2016). The histogram representation is, however, inconvenient, because the height of the histogram depends on the width of the size bin that are used. This led Platt and Denman (1977) to introduce the “normalized size spectrum” as the biomass distribution as a function of cell size *b*(*m*). Being a distribution means that the spectrum has dimensions of biomass per cell mass, and that the integral of the spectrum is the total biomass. It is convenient to introduce the “Sheldon spectrum” as *b*(*m*)*m* because it has the same property as the log-binned histogram that it is approximately flat (Box 4).

The flat Sheldon spectrum is commonly understood as emerging from predator-prey interactions. First, Sheldon et al. (1972) showed how the biomass in successive trophic levels scaled as *E*_T_*β*^0.25^ ≈ 0.9, where *E*_T_ ≈ 0.2 is the trophic efficiency and *β* ≈ is the predator prey mass ratio. This results was later re-derived as part of the metabolic theory of ecology (Brown et al., 2004), however, only by introducing an extra assumption about energy equivalence. The result relies on the trophic efficiency, which is a quantity that is hard to estimate, and which eventually is an emergent property of the community structure (Borgmann, 1987). An alternative argument by Andersen and Beyer (2006) derived the size spectrum purely based on individual-level properties. As all of these arguments only rely upon predator-prey interactions, it is not clear how well they apply among the lower trophic levels of the ocean where many cells mainly subsist on photosynthesis and recycled production from dissolved organic matter and less on predation on smaller particles. Poulin and Franks (2010) refined the argument by considering phytoplankton and zooplankton spectra separately to show a flat phytoplankton spectrum and a declining zooplankton spectrum. Here we will explain the scaling of the community size spectrum only from considerations of predator-prey interactions by an extension of the Andersen and Beyer (2006) argument, and later show that the predictions fit surprisingly well with dynamical simulations.

Predator-prey interactions are described by bigger cells predating on smaller cells (Hansen et al., 1997). The size preference for predation can be described by a log-normal size selection function:

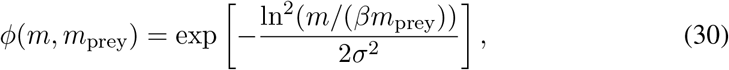

where *m*_prey_ is prey size, *β* the preferred predator:prey mass ratio and *σ* the width of the preference function. The available food is found by integrating across all size groups:

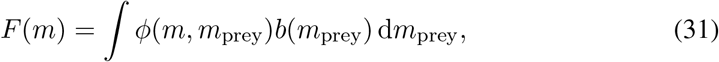

where *b*(*m*) is the biomass size spectrum. From this description we can derive the size spectrum as (Box 5):

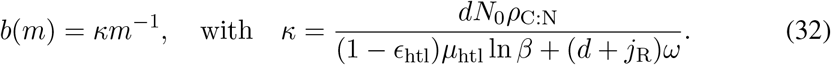

The spectrum scales with mass as *m^−^*^1^, which means that the Sheldon spectrum *∝ b*(*m*)*m* is constant. The height of the spectrum (the coefficient *κ*) is a novel result. The height depends on the mixing rate *d*, the concentration of nutrients being mixed up from the deep *N*_0_, the mortality imposed by higher trophic levels *μ*_htl_, and the length of the size spectrum *ω* = ln(*m*_max_/*m*_min_). The main controlling parameter is the mixing rate. The height of the spectrum increases with mixing but saturates at high mixing rates (*d ≪ μ*_htl_ ln *β/ω*).

At very high mixing rates the production will be limited by the synthesis capacity of the cell, which is not accounted for here, however, that is probably a rare occurrence in nature.

### 4.5 Dynamic size-based model

Further insight into the size structure requires numerical simulations. Here we simulate the entire unicellular plankton community by embedding the model of cell resource uptake and metabolism in a simple ecosystem model. Cells are divided into size group with each group *i* representing the biomass *B_i_* within a range of cells with the geometric mean mass *m_i_*. For simplicity we have assumed that cells have constant C:N mass ratio *ρ*_C:N_ = 5.68, but the model can be extended to dynamic stoichiometry (Ho et al., 2020; Ward et al., 2018). The rate of change (the growth rate) of biomass in a size group is:

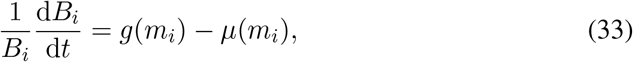

where *g*(*m*) is the division rate and *μ*(*m*) is the total morality. The division rate is determined by resource encounter and synthesis (Table 4, eq. M4).

**Table 4:** Processes and equations to calculate the division rate *g* of a cell (note that the population growth rate also requires the subtraction of losses). All rates are in units of g_C_/g_C_ per time.

|  |  |  |
| --- | --- | --- |
| *Encounter and synthesis:* |  |  |
| Available carbon | $j_{C.net} = j_{DOC} + j_L + j_F - j_R - j_{passive}$ | (M1) |
| Available nutrients | $j_{N.net} = j_N + j_F - j_{passive}$ | (M2) |
| Leibig's law | $j_{net} = \min\{j_{C.net}, j_{N.net}\}$ | (M3). |
| Synthesis | $g = j_{max} \frac{j_{net}}{j_{net} + j_{max}}$ | (M4) |
| *Down-regulation of uptakes* |  |  |
| Feeding | $\tilde{j}_F = \max\{0, j_F - (j_{net} - g)\}$ | (M5) |
| Photoharvesting | $\tilde{j}_L = j_L - \max\{0, \min\{(j_{C.net} - (j_F - \tilde{j}_F) - g), j_L\}\}$ | (M6) |
| DOC uptake | $\tilde{j}_{DOC} = g - \tilde{j}_F - \tilde{j}_L$ | (M7) |
| *Mortalities:* |  |  |
| Predation: | $\mu_p(m_j) = \sum_i \frac{\tilde{j}_{F,i}}{\epsilon_F} \frac{\Phi_{ij}}{F_i} B_i$ | (M7) |
| Viral lysis: | $\mu_{v,i} = \mu_{v0} B_i$ | (M8) |
| Higher trophic levels: | $\mu_{htl}(m) = \begin{cases} \mu_{htl0} \phi(m, m) & \text{for } m < m_{htl} \\ \mu_{htl0} & \text{for } m \geq m_{htl} \end{cases}$ | (M9) |
| All fluxes are calculated according to the relations in Section 3. Available food is $F_i = \sum_j \Phi_{ij} B_j$ where the effective preference between size groups $\Phi_{ij}$ is found by integration across the width of the size groups (Appendix A). The tildes above the uptakes of light and food indicates down-regulation in eqs. (M5-6). (M1-2): Uptakes are given by Eq. 3 combined with affinities for nutrients and DOC (Eq. 5), light (Eq. 13) and food (Eqs. 15 and 31). (M3): a standard functional type II response aka. ‘‘Monod’’ function. (M7): The predation mortality exerted by unicellular plankton can be calculated as the ratio between the amount of food eaten by all predators from size group $j$ and biomass at size group $j$. The amount eaten is: $E_j = \sum_i (\tilde{j}_{F,i}/\epsilon_F) B_i \phi(m_i, m_j) B_j / F_i$. Moving $B_j$ outside the sum to calculate predation mortality as $\mu_{p,j} = E_j / B_j$ gives M7. |  |  |

Mortality has three origins: predation by unicellular plankton *μ*_p_ through the process of big cells eating smaller cells (M7), viral lysis *μ*_2_ (M8), and predation by higher trophic levels *μ*_htl_ (M9). Viral mortality is modelled by assuming that viral mortality is proportional to biomass. Mortality by higher trophic levels acts on the largest size groups. We use a selection function consisting of combination of the logarithmic size selection function in Eq. 30 and a constant level.

Nutrients and DOC are updated with the uptakes and losses from the cell-level processes:

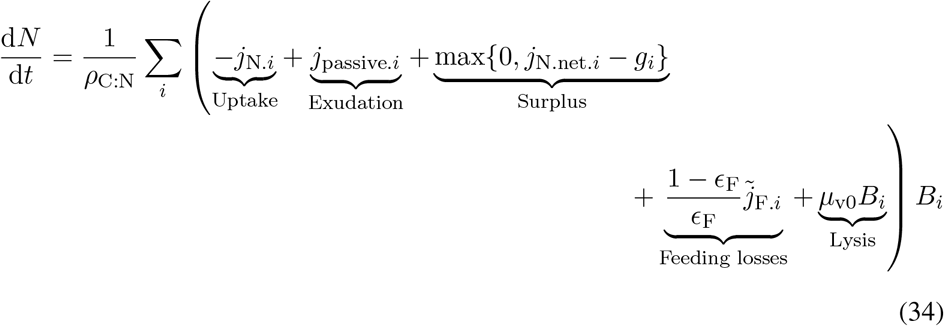

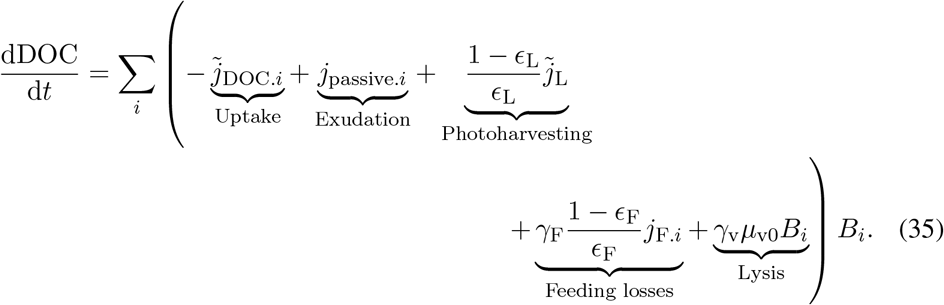

Generation of *N* happens through assimilation losses, passive exudation and remineralised viral lysis. Generation of labile DOC happens through passive exudation, assimilation losses from light harvesting and phagotrophy, and from remineralized viral lysis. We assume that all nutrient losses from viral lysis are made available over short time scales (Carlson, 2002), but only a fraction *γ*_v_ of carbon losses are labile. All the losses to higher trophic levels are eventually converted to particulate organic matter (which is not explicitly resolved here) so there is no remineralization of those losses.

Five parameters control the chemostat: mixing rate *d*, deep nutrient concentration *N*_0_, light *L*, temperature *T* and the mortality imposed by higher trophic levels *μ*_htl0_, however, only three are important. The mixing rate and the deep nutrient concentration mainly enter as a product so we can focus on only one of them – the mixing rate is commonly chosen. In a water column the productive layer will adjust itself to the depth where cells are co-limited by light and nutrients (Ryabov et al., 2010; Beckmann and Hense, 2007; Klausmeier and Litchman, 2001). The light level is therefore also of minor importance, as long as it is sufficiently high to not be limiting. We first concentrate on the mixing rate *d* and take up the importance of higher trophic level mortality and temperature in the follow section.

Chemostat simulations in eutrophic situations with a high mixing rate show an extended flat Sheldon spectrum occupying the full size range (Fig. 9). The level of the size spectrum fits with the theoretical prediction (Eq. 27). Phototrophic cells span a wide size range and due to the high influx of nutrients they are light limited and not nutrient limited. Microplankton and partly nanoplankton have a significant influx of carbon and nutrients from phagotrophy. Only the largest cells are fully heterotrophic in the sense that they leak surplus nutrients from phagotrophic uptakes.

**Figure 9:**
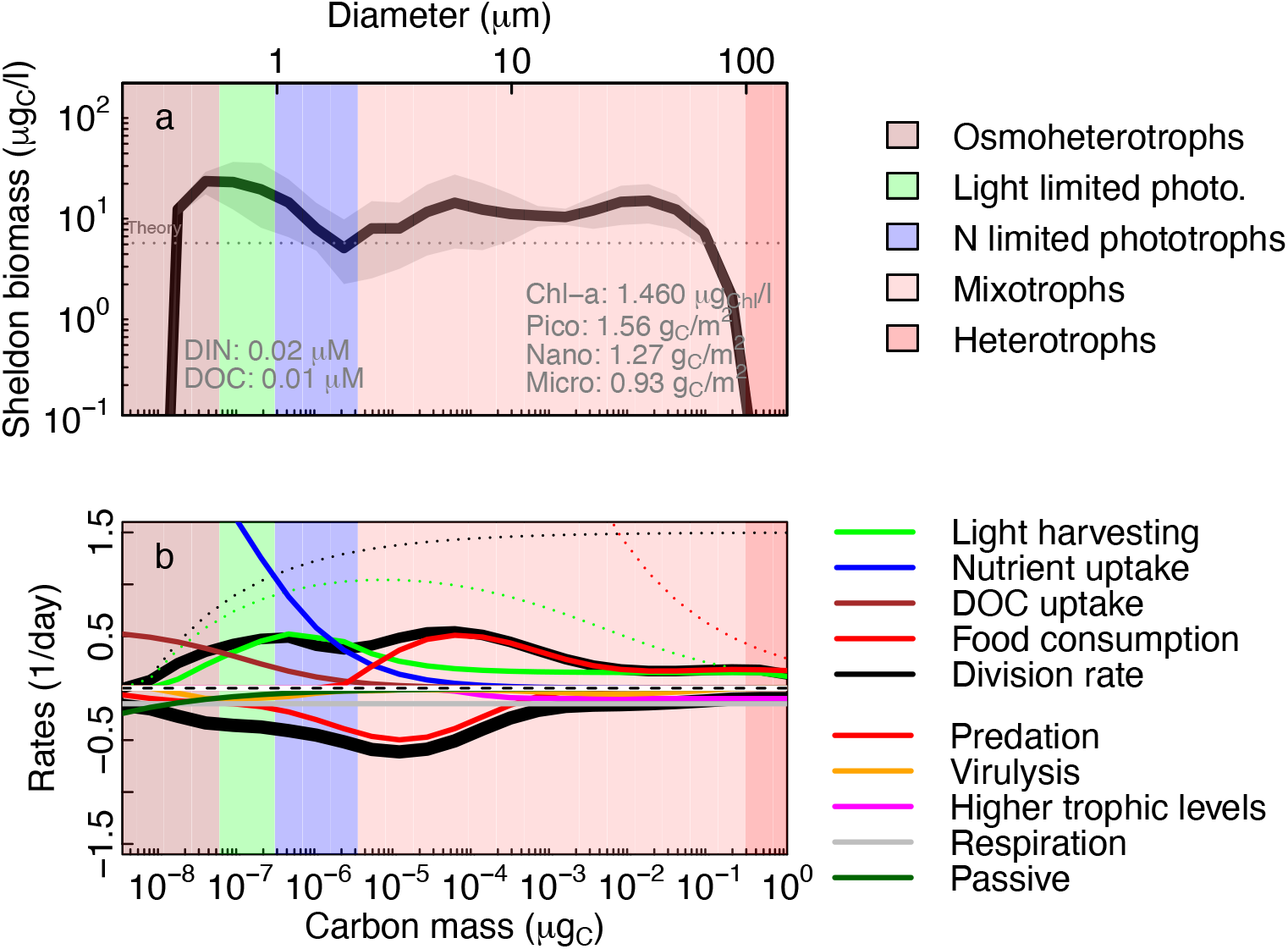
Results of simulations under eutrophic conditions with high mixing. a) Sheldon size spectrum (Box 5). The background colours indicate the trophic strategy (see Fig. 8). b) Rates of uptakes and losses in biomass specific units (day ^−1^). The dotted lines show maximum possible uptakes or growth rates. The thick black lines in panel b are the total division and loss rates (note that in this case they are not equal as the simulation is not in a steady state; the variation is indicated in panel a with the grey area around the mean). Parameters: Light *L* = 40 μE/m^2^/s, mixing rate *d* = 0.1 m/day, and higher trophic level mortality *μ*_htl_ = 0.1 day ^−1^.

**Figure 10:**
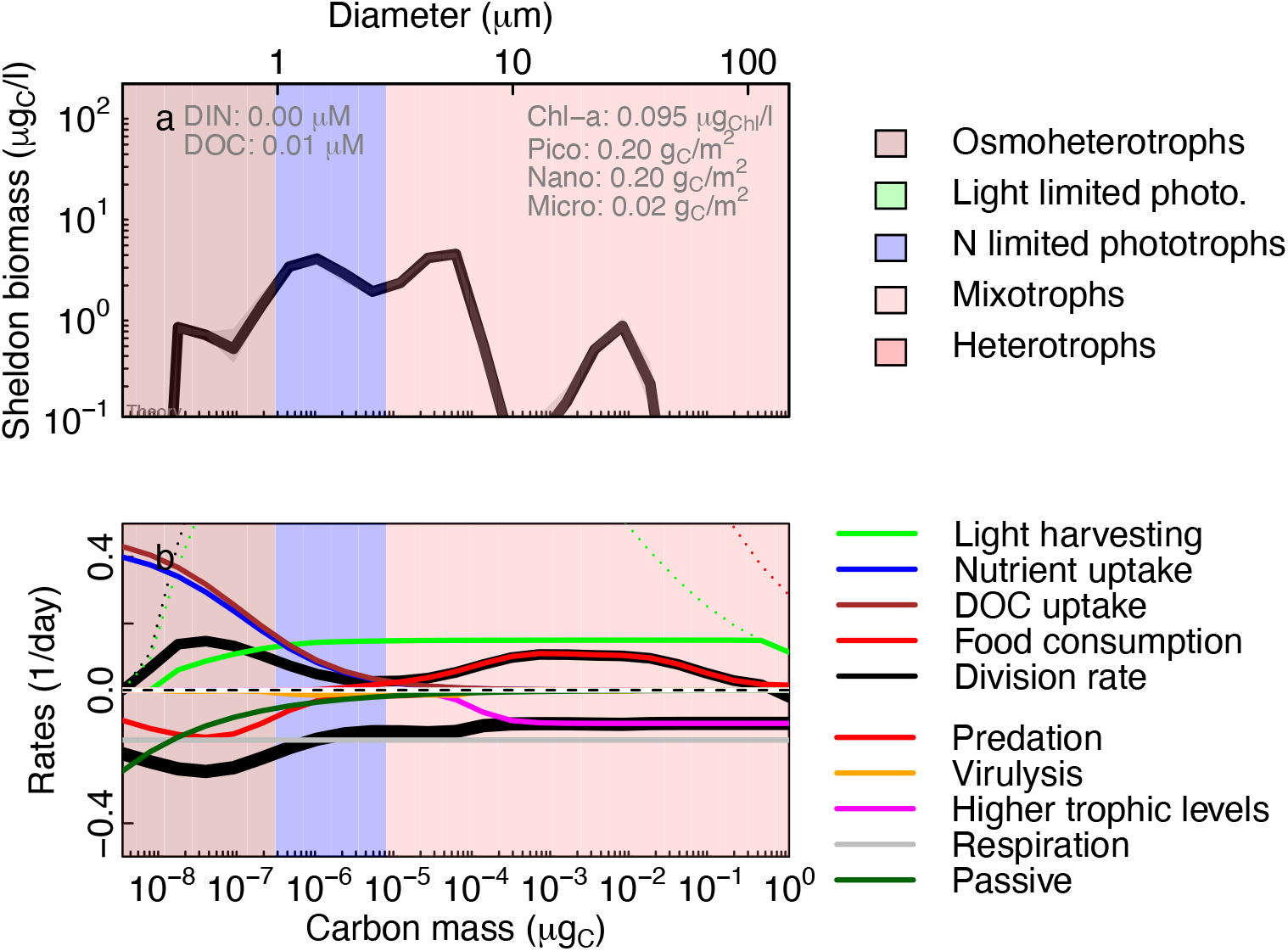
Results of simulations for an oligotrophic situation with low diffusivity of nutrients; see Fig. 9 for explanation (note different y-axis on panel b). Mixing rate *d* = 0.001 1/day.

Under oligotrophic situations with a low mixing rate the spectrum is still flat, but the realized size range is smaller in both ends of the spectrum. Therefore the “height” of the spectrum (*κ*) is also higher than predicted, because the prediction was only valid for a full range spectrum. Under oligotrophic situations the phototrophs are fully nutrient limited. It is unrealistic that very small cells are absent in oligotrophic conditions, because they are expected to dominate. This is because of the rising limiting nutrient values (*N ^*^*) of small cells due to the limitation imposed by the cell membrane (Fig. 7).

Simulating across mixing rates shows generally flat Sheldon size spectra (Fig. 11). The spectrum exponent (panel e) is roughly zero and the size spectrum level (*κ*) follows the prediction from Eq. 27 (panel d). At small mixing rates small picoplankton dominates. As mixing rate increase, the upper size range extends. The extension of the upper size increases the length of the food chain. This result was demonstrated in a simple size-based by Armstrong (1994) and confirmed by alternative derivations by Poulin and Franks (2010) and Ward et al. (2014). At very high mixing rates the spectrum again becomes truncated. The truncation at high mixing rates is a result of plankton being mixed out of the productive layer faster than the maximum growth rate. Overall it is clear that the overall size spectrum exponent is unaffected by the environmental conditions, only the height and the extent of the spectrum are affected.

**Figure 11:**
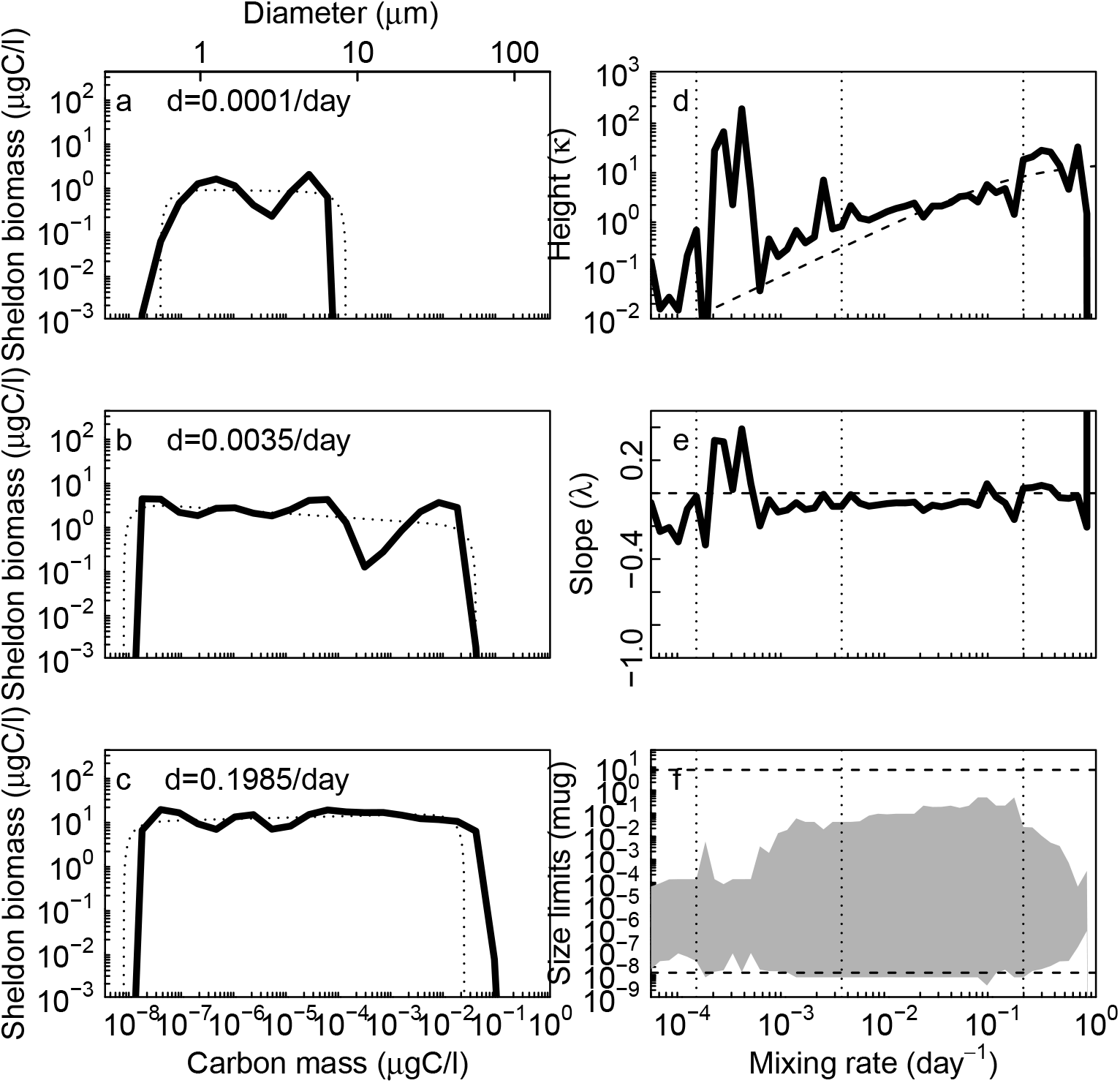
Size spectra with varying mixing rates under high light. Panels a-c: size spectra fitted to a power-law (Eq. 32) truncated at high and low sizes (dotted). Panels d-f: results of the fits for height (*κ*), exponent (*λ*) and upper and lower size limits. The dashed lines are the theoretical predictions of *κ* from Eq. 27, exponent *λ* from Eq. 32, and min and max sizes from Eqs. 20 and 21. The “wiggles” are due to inaccuracies in the fit of the size spectra. The vertical dotted lines in panels d-f show the three mixing rates used to calculate panels a-c. *L* = 100 μE/m^2^/s.

## 5 Ecosystem functions

The size distribution combined with the cell-level characteristics allows the calculation of ecosystem functions. Ecosystem functions can be divided into biomasses, production, and efficiencies. Because size-based models consider cells with multiple trophic strategies, calculating the functions are somewhat different than for ordinary functional group type of models (see Table 5). In the chemostat model the integration over the water column comes about simply by multiplying with the thickness of the productive layer that we arbitrarily set to *M* ≈ 20 m.

**Table 5:** Ecosystem biomass and functions. *M* is the thickness of the mixed layer, here set to 20 m.

|  |  | Value |  |  |
| --- | --- | --- | --- | --- |
| Function | Formula | Olig. | Eut. |  |
| *Biomasses* |  |  |  |  |
| Biomass | $B_{\text{total}} = M \sum_i B_i$ | 0.38 | 3.8 | $\text{g}_\text{C}/\text{m}^2$ |
| Chlorophyl<sup>(1)</sup> | $B_{\text{Chl}} = M \sum_i \tilde{j}_{\text{L},i} B_i / L$ | 0.085 | 1.5 | $\mu\text{g}_{\text{Chl}}/\text{l}$ |
| *Production* |  |  |  |  |
| Gross PP | $P_{\text{gross}} = M \sum_i \tilde{j}_{\text{L},i} B_i / \epsilon_{\text{L}}$ | 20 | 360 | $\text{g}_\text{C}/\text{m}^2/\text{yr}$ |
| Net PP | $P_{\text{net}} = M \sum_i \max\{0, \tilde{j}_{\text{L},i} - j_{\text{R},i}\} B_i$ | 0 | 150 | $\text{g}_\text{C}/\text{m}^2/\text{yr}$ |
| Net bact. prod. | $P_{\text{bact}} = M \sum_i \max\{0, j_{\text{DOC},i} - j_{\text{R},i}\} B_i$ | 0.12 | 66 | $\text{g}_\text{C}/\text{m}^2/\text{yr}$ |
| New prod. | $P_{\text{new}} = M d(N_0 - N) \rho_{\text{C:N}}$ | 2.1 | 210 | $\text{g}_\text{C}/\text{m}^2/\text{yr}$ |
| Prod. of HTLs | $P_{\text{htl}} = M \sum_i \mu_{\text{htl},i} B_i$ | 2.5 | 60 | $\text{g}_\text{C}/\text{m}^2/\text{yr}$ |
| *Efficiencies* |  |  |  |  |
| Eff. of PP | $\epsilon_{\text{PP}} = P_{\text{net}} / P_{\text{gross}}$ | 0.0 | 0.43 | |
| Eff. of bact. | $\epsilon_{\text{bact}} = P_{\text{bact}} / P_{\text{net}}$ | - | 0.42 | |
| Eff. of HTL | $\epsilon_{\text{htl}} = P_{\text{htl}} / P_{\text{net}}$ | - | 0.43 | |
| <sup>(1)</sup> Mass of Chl per carbon mass is approximately proportional to the down-regulated |  |  |  |  |
| mass-specific light affinity $\tilde{j}_{\text{L},i} / L$. (Edwards et al., 2015). | | | | |

Gross primary production is the total amount of carbon fixed:

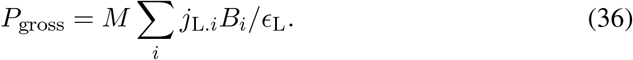

The net production is the carbon available for biomass production, i.e., the gross primary production minus the exudation losses and respiration. That definition, however, only works for purely phototrophic plankton. Here, plankton are mixotrophs and some larger mixotrophs may contribute negatively to primary production because they fix negligible amounts of carbon. To compensate, we consider only the groups where net fixation (fixation minus respiration) is positive. This procedure assumes that carbon from photosynthesis is prioritized for respiration over other carbon sources (DOC or feeding). Bacterial production is net production based on DOC uptake. It faces a similar problem as the net primary production, and again we only consider positive net contributions. New production is the amount of nutrient that diffuses up into the photic zone to fuel primary production. Finally, we can calculate the production to higher trophic levels as the losses to higher trophic level mortality.

Efficiencies the ratio between a production and the net primary production. They are typically in the range 0 to 1.

The total biomass is roughly proportional to the mixing rate *d* (Eq. 27) until it becomes limited by light, around *d* = 0.02 day^−1^ in low light conditions (Fig. 12 a+f). The total Chl-concentration varies between 0.01 to 1 μgChl/l (panels b+g), which is in line with outputs of global circulation model simulations (Van Oostende et al., 2018). When production becomes light limited the nutrient level increases because plankton production cannot fix all the available nutrients (at mixing rates of 0.01 and 0.1 day^−1^ in low and high light respectively). The increases in biomass is reflected in the productions, which also increase roughly proportional to the biomass (panels c+h). However, the relative magnitude of the different productions changes with mixing rate, as reflected in the efficiencies (panels d+i). The production efficiency generally increases with the mixing rate. Surprisingly, the higher trophic level production can become larger than net primary production resulting in *∊*_htl_ > 1. This occurs in oligotrophic situations where a high gross primary production fuels a high DOC production from exudation. That DOC also fuels plankton production, which eventually manifests itself in high net production to higher trophic levels. The gross efficiency of higher trophic level production (*P*_htl_/*P*_gross_ will always be < 1.

**Figure 12:**
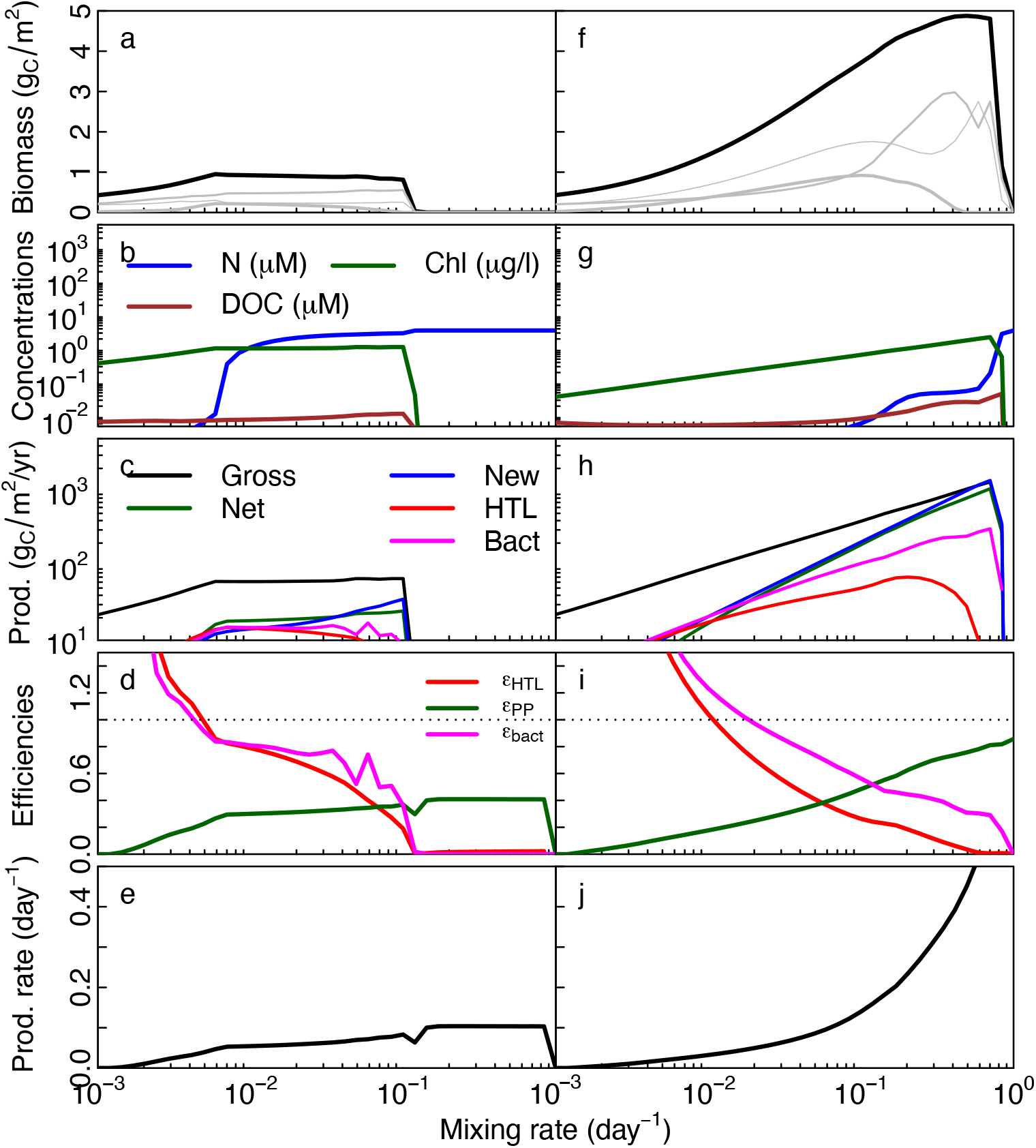
Ecosystem biomass and functions at low and high light (*L* = 10 and 100 μmol m ^−2^s ^−1^). In the top row the gray lines show pico-, nano-, and microplankton. The last row shows the community production rate as the net primary production divided by biomass.

### 5.1 DOC and bacteria production

A difficult aspect of the ecosystem model is to parameterize the production and uptake of dissolved organic matter (DOC). Part of the difficulty stems from our incomplete knowledge of DOC: how much is labile and how much is not? And further: what are the sources of DOC: how much DOC is produced by incomplete assimilation and how much by passive exudation – or between “income” and “property” taxes (Bjørnsen, 1988)? In the ecosystem model DOC represents the labile DOC that can be immediately taken up and used. Labile DOC is produced by incomplete assimilation of photoharvesting (*E*_L_), from passive exudation (*j*_passive_), from assimilation losses due to feeding (*E*_F_), and from viral lysis (*j*_v_).

Pelagic ecosystem models typically describe DOC release as a constant fraction of fraction of primary production (Thornton, 2014), though some include size-based passive exudation (Kriest and Oschlies, 2007) using Bjørnsen (1988)’s model, or a more complex division between labile and non-labile pools (Anderson and Williams, 1998; Flynn et al., 2008). The size-based model represents all processes: passive size-based exudation, exudation due to uptake, incomplete feeding, and viral lysis, but does not distinguish between labile and refractory DOC.

All the incompletely assimilated carbon from photoharvesting is assumed to be available as DOC. The assimilation fraction is commonly set between 2-10%; following Anderson and Williams (1998) we use 20% (*E*_L_ = 0.8) to have sufficient DOC available.

Passive losses are discussed in section 3.2; we assume that all passive losses are labile.

Feeding losses from phagotrophy are set to 20% (*E*_F_ = 0.8). However, not all feeding losses may be available, as mostly non-digestible material will be exuded; Anderson and Williams (1998) send only 10% of feeding losses back to DOC so we set the available fraction at *γ*_F_ = 0.1. There is disagreement about the fraction of lysed cells that is available as labile DOC. Carlson (2002) find that the majority of the dissolved organic matter released from bacterial lysis is available, while (Anderson and Williams, 1998) only assume that 3.4% is available as labile DOC. We assume that half is available; *γ*_v_ = 0.5. Feeding by higher trophic levels could also lead to DOC production. We assume that most sloppy feeding by higher trophic levels lead to particulate organic matter, which is not represented here, so *γ*_htl_ = 0.

The model gives total DOC losses around 30 % (Fig. 13c+d), which is within the expected range. Overall we expect the average losses to DOC from passive exudation and assimilation losses from feeding to be 10-30 % of their production (growth rate) (Kiørboe, 1993; Carlson, 2002). This is within the range simulated by the model Fig. 13a+b. Smaller cells have much higher passive losses, which is essentially the process that determines the lower cell size.

**Figure 13:**
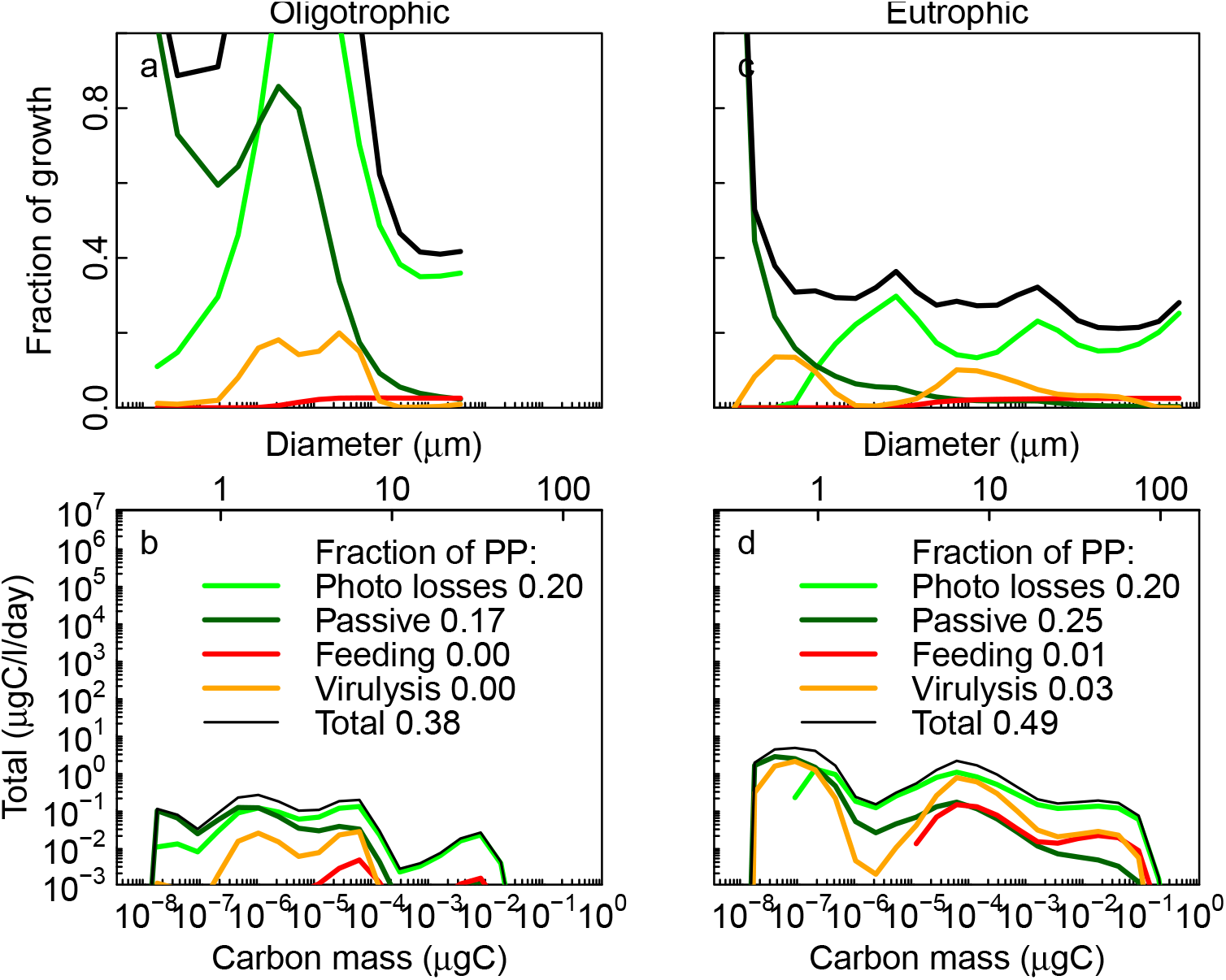
Losses to DOC as a function of cell size for oligotrophic conditions (Fig. 4.5) and eutrophic conditions (Fig. 9). Top row: losses as fraction of cell growth rate. Bottom row: total losses with fractions of gross primary production given in the legend.

The fraction of primary production becoming labile DOC should vary between 2-40%; highest in oligotrophic waters (Teira et al., 2001). In productive regions DOC originates mainly from passive exudation while assimilation losses from feeding supply DOC in oligotrophic regions (Teira et al., 2001). The observed fraction of primary production exuded is 2-40 % (Teira et al., 2001; López-Sandoval et al., 2013); highest in oligotrophic regions. The model have a total losses in the same range (Fig. 13c+d legend). An important source in eutrophic waters is viral lysis while in oligotrophic waters the main source is photoharvesting.

Regarding the size-scaling of DOC losses the evidence is conflicted. Kiørboe et al. (1990) sees strong evidence of size-scaling of passive exudation, Teira et al. (2001) sees some indirect evidence, while Maranón et al. (2004) did not see any evidence (but notes that nutrient limitation may be a confounding factor). The diverging evidence reflects the difficulty in distinguishing between different sources of DOC and that studies focus on different size-ranges of cells. For example, López-Sandoval et al. (2013) notes that there is no overall size-scaling of DOC exudation among the plankton, which may be due to different processes dominating among small cells (passive exudation) and large cells (assimilation losses). The modelled total amount of DOC losses is roughly independent of size (Fig. 13b+c), though with higher passive losses for small cells. Among smaller cells the main source of losses are a combination of passive exudation, photosynthesis losses, and viral lysis. Among larger cells feeding assimilation losses are a potential important term, which is, however, limited by the assumption that only a small fraction of feeding losses are labile (*γ*_F_ = 0.1).

The only quantitative evidence on the size-relation between bacterial production and cell size is that the bacterial generation time is inversely correlated with the total surface area of plankton cells with a power-law exponent −0.82 (Kiørboe et al., 1990). The model gives a similar relation (Fig. 14), though with slower generation times. This may have to do with how the average generation time is calculated. In the model the generation time also includes cells with very slow DOC uptake rates, which increased the average generation time.

**Figure 14:**
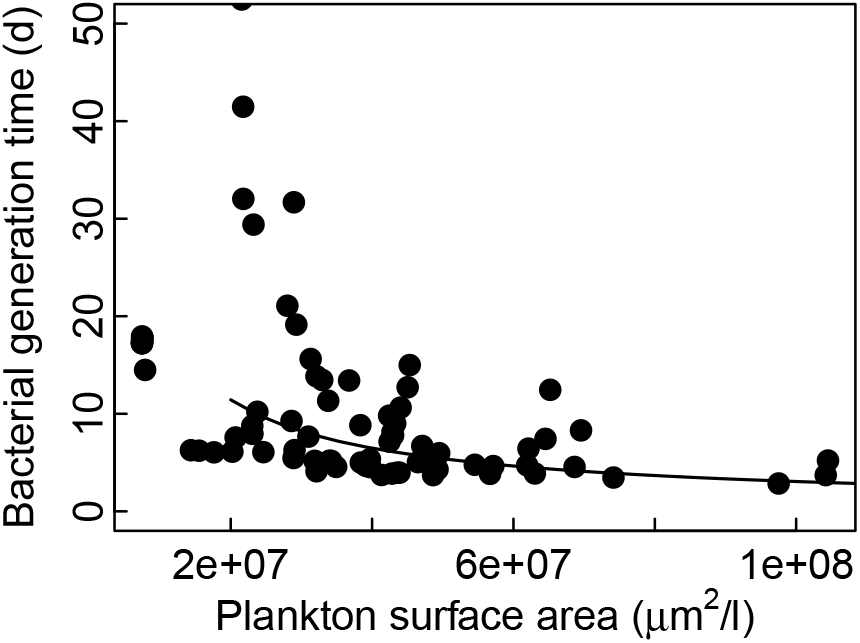
Bacterial generation time as a function of the total surface area of plankton in the size range ESD = 5 to 60 μm. The bacterial productivity is calculated as the flux of DOC uptake (*j*_DOC_) minus losses to passive exudation and respiration: max {0, *j*_DOC_ — *j*_passive_ — *j*_R_}. The generation time is 1 divided by the average of all bacterial productivities larger than 0. The lines shows a fit to a power law with fixed exponent 0.82. The mixing rate ranges between 0.001 and 0.1 d ^−1^ and light is from 10 to 60 μE/m^2^/s.

The modelled concentrations of labile DOC are very low (around 1 *μ*M; Fig. 12b+g). This is because all DOC is considered labile and it is therefore immediately taken up and drawn down towards limiting concentrations which are around 0.1 *μ*M (Fig. 7). Including also refractory DOC would allow for higher DOC concentrations.

### 5.2 Effect of higher trophic level mortality

The model results depend upon the mortality exerted by the larger multicellular organisms as represented by the higher trophic level mortality *μ*_htl_. The importance of the HTL mortality is not unique to size-based models; results of all plankton model are sensitive to this closure term. However, the important effects of this closure term is rarely acknowledged (but see Steele and Henderson, 1992). Varying the HTL mortality affects the size structure and the functions of the plankton ecosystem (Fig. 15). The main effect of increasing higher trophic level mortality is to truncate the size-spectrum. The truncation releases the smaller plankton from predation and they respond by becoming more abundant. Due to this trophic cascade the total biomass and the net primary production are only weakly affected by the HTL mortality (Fig. 15b). The main effect is on the production towards the higher trophic levels, which has a uni-modal shape with a maximum at intermediate mortalities. This behaviour is similar to how fisheries models responds to fishing where the maximum is termed the “maximum sustainable yield”. In lower productive system the effect of HTL mortality is stronger and the peak in HTL mortality is reached at lower mortalities (not shown).

**Figure 15:**
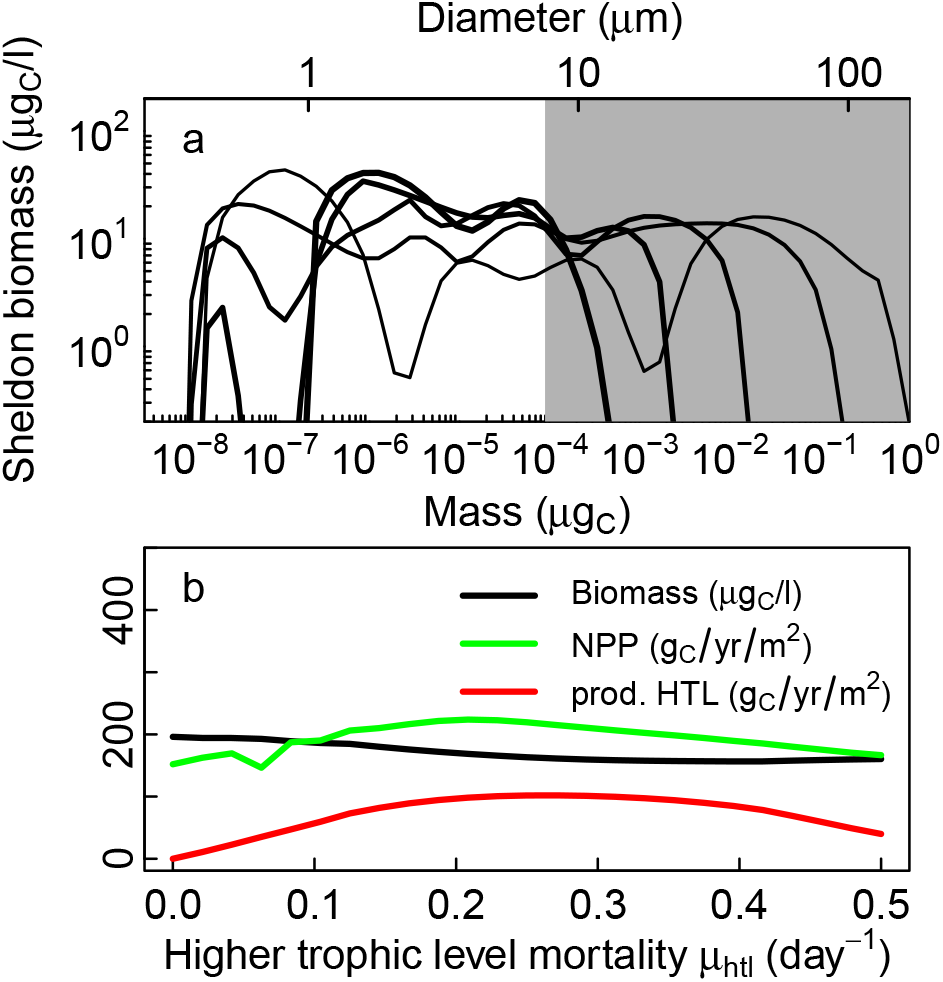
Effect of varying the mortality exerted by higher trophic levels. a) Sheldon size spectra for higher trophic level mortality from 0 to 0.5 day ^−1^ (thin to thick lines). The grey patch illustrates the size-range where HTL mortality acts directly. b) Effect of HTL mortality on ecosystem functions: total biomass, net primary production, and HTL productivity. Light and mixing are as the eutrophic situation Fig. 9

Since the HTL mortality is an extrinsic parameter, we would like to know a reasonable value. That is difficult because the level of mortality depends on the predators: higher productivity of the larger plankton will lead to a higher HTL mortality. This effect is also seen inside the spectrum on the level of the predation mortality in Figs. 4.5-9b: in the oligotrophic system the level of mortality on the smallest plankton is around 0.1 day^−1^rising to around 0.25 day^−1^in the eutrophic system. HTL mortality should therefore not be higher than 0.25 day^−1^. Here we have used 0.1 day^−1^.

### 5.3 Effects of temperature

Water temperature directly affects affinities and metabolism and, through this, a host of processes from the division rate of cells, to ecosystem structure and functions (see Section 3.4; Fig. 16). Higher temperature increases the division rates of all cell sizes up to a point where division rates begin to decrease (panels a+d). The increase is relatively modest, though, and much less than indicated by a “metabolic” *Q*_10_ = 2, even for large heterotrophic cells. It is also less than the *Q*_10_ values often used in plankton simulation models (e.g. Archibald et al., 2022). The reason for the relatively slow increase in division rates is that the cells are generally limited by encounter with resources (nutrients, light, and prey), which has a small *Q*_10_. The decrease at higher temperature occurs when respiration losses, with a high *Q*_10_, begins to dominate over the resource uptakes with smaller *Q*_10_’s. The increasing division rates have a modest effect on ecosystem functions and structure (panel b-e), but they do increase net primary production and increases the maximum size in the size spectrum. The temperature response is, however, very dependent upon the conditions. Under oligotrophic conditions the temperature response is almost absent. The oligotrophic situation is dominated by carbon input from phototrophy (Fig. 9), which is independent of temperature (*Q*_10_ = 1). Furthermore, light is available in excess (dotted green line in Fig. 9b) and can easily support the basal metabolism. Clearly, the response to temperature depend on the mixing rate and light in complicated ways (Serra-Pompei et al., 2019) making it hard to make generalisations.

**Figure 16:**
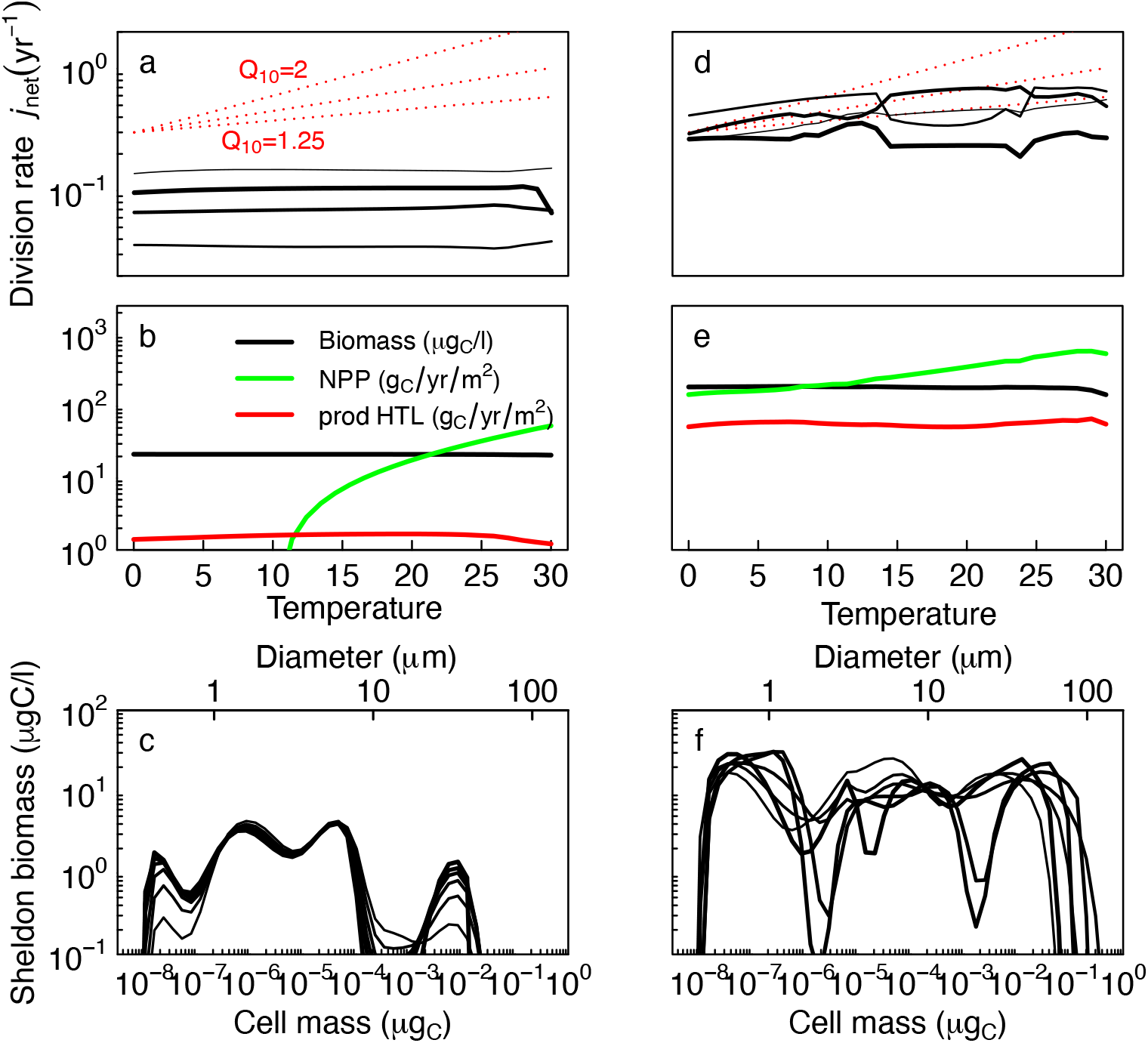
Effect of varying the temperature between 0 and 25 degrees in oligotrophic/eutrophic conditions (left/right columns; Fig. 9). a,d) Effect on the division rate *j*_net_ for 4 size classes (increasing line width). The red lines show temperature increases described by *Q*_10_ of 1.25, 1.5 and 2. b,e) Effect on ecosystem functions: total biomass, net primary production, and HTL productivity. c,f) spectra for 6 temperatures (increasing line widths).

## 6 Discussion

We have reviewed how cell-level processes can be related to cell size and first principles, and how they ultimately determine major aspects of plankton community structure and function. The approach builds upon the central role of cell size for resource uptake and metabolism of unicellular plankton. By cultivating a view of each cell as a “generalist” that can perform all types of resource uptakes – essentially being a combination of a bacteria, a phytoplankton, and a zooplankton – the trophic strategies become an emergent property. The fundamental processes at the cell level is based upon existing theory and knowledge with a few novel elements: a fluid-mechanical argument for the clearance rate, the upper limit of phagotrophic assimilation, and the identification of two scaling regimes for light affinity. The synthesis of all processes in a dynamic “minimal” size-based, along the lines of Ward and Follows (2016) and Andersen et al. (2015), leads to a complete ecosystem model that resolves the community size spectrum as well as the dominant trophic roles of plankton of different sizes. Novelties in the minimal model is the inclusion of dissolved organic carbon to represent carbon reuse and the microbial loop, and the development of closed-form analytical solutions for the scaling and level of the size-spectrum. Throughout we have maintained a focus on simplicity of all processes to bring forth a clear understanding of how each process contributes to the community structure. Despite that operational plankton models – even those only based on cell size – are more complex and complete than the minimal framework analysed here, the effects of nutrient enrichment, higher trophic level mortality, temperature, and light upon the structure and function of the community are likely to be universally present.

The model generally reproduce observed ranges of biomass, chlorophyll, and productivity as observed in natural systems. The structure of the ecosystem is determined by a combination of the bottom-up processes from nutrient availability and light, by the internal process of predation, and by the top-down process of higher trophic level predation. As also shown by Poulin and Franks (2010) the availability of nutrients determine the potential length of the food chain (the maximum size). This result is similar to the classic insight in theoretical ecology about resource productivity determining food chain length (Oksanen et al., 1981). However, the top-down effect of higher trophic level mortality plays a key role in the structure and function of the community (Steele and Henderson, 1992). From a modelling perspective, this is problematic, as it to some degree ruins the universal nature of the model: in a given situation the level of the higher trophic level mortality needs to be determined. In global simulations the higher trophic level mortality will vary depending upon the predation pressure from the multicellular plankton community, which is not the same throughout the global ocean. One solution is to use a higher trophic level mortality that varies linearly with plankton biomass (a “quadratic” loss term); a more complex solution is to include a representation of multicellular plankton (Serra-Pompei et al., 2020). In terms of size spectrum slope, the model generally reproduce the commonly observed flat Sheldon spectrum (Sprules and Barth, 2016; Kenitz et al., 2019). The model therefore reproduces the conclusion by Poulin and Franks (2010) that the spectrum slope in itself is not informative of the plankton structure.

In the following we discuss how first principles constrain the cell’s function and which processes are still weakly constrained. Second, we discuss the limitations of the size-based approach to modelling plankton communities and how it relates to trait-based and functional-group based plankton models.

### 6.1 Parameters from first principles or empirical meta analyses

“First principles” are relations rooted in physics, chemistry, evolution, or geometry. Tying descriptions of processes and parameters to first principles has succeeded to varying degrees. We distinguish between four levels of success: *i*) The process is known and the parameter(s) can be calculated from first principles; *ii*) scaling exponents with cells size (mass, volume, or diameter) are known from first principles but the coefficients have to be calibrated with laboratory measurements or from meta-analyses; *iii*) The governing process is known but the theoretical argument has not been developed and parameters rely solely on empirical knowledge; *iv*) The empirical evidence is lacking and parameters are only constrained indirectly via loose arguments or tuning of the outcome with observations of the community structure. For unicellular plankton all four levels are encountered.

The most complete level (*i*) description is how diffusion limits uptake of dissolved nutrient or DOC uptake and phototrophy, including the temperature scaling. For phototrophy there are clear theoretical arguments for all coefficients and scaling exponents – including the novel argument for the transition between two scaling regimes – simply from geometric arguments. However, a better understanding of the quantum yield is needed to fully describe the observed variation in light affinity. The other well known effect is how the cell membrane limits the lower size of a cell, though the thickness of the cell wall is not (yet) constrained by first principle arguments. Another example of the role of geometry, which is not explored here, is the importance of a vacuole, a principle characteristic of diatoms, to modify the diffusion uptake (Hansen and Visser, 2019; Cadier et al., 2020). The other level (*i*) description is the novel argument of how clearance rate of prey encounter is derived from fluid mechanics, though it must be recognised that the amount of energy available to the cell is a guesstimate. As the fraction of energy only enters as a square root in Box 3 this value is not crucial. Comparing the theoretical result with data (Fig. 4) shows a scatter of *±* 1 order of magnitude. It is known how the variation in clearance rate across the mean is due to the hydrodynamics of different flagella arrangements that also results in different predation risk (lower clearance leads to smaller predation risk; Nielsen and Kiørboe (2021)) – we return to this in the next section.

Most processes belong to the second level where scaling exponents are well described but empirical knowledge is needed to determine exact parameter values. To this category belong the processes related to the “secondary” scalings of nutrient and light uptake, i.e., that the scaling is flat for small sizes. We can confidently argue that the scaling should be flat, but cannot determine the value of the parameters (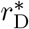 and 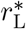). From the simulations we see that these flat scalings do not have a strong impact on the resulting ecosystem, so one could even omit them without a great loss of accuracy. Other level two processes are the passive exudation, metabolism, and the temperature scaling of metabolism. The temperature dependence of metabolism is a complicated mixture of many different processes and a simple first-principles argument does not exist.

Finally, some of the parameters associated to DOC losses are largely guesswork (level iv). The poor state of knowledge is partly due to our limited understanding of the enormous diversity of DOC compounds and their lability, which makes the lumping of DOC into one group crude. We note that this problem is recently receiving attention (Zakem et al., 2021) and hope that a better understanding of DOC is forthcoming.

Overall, while it is clear that first principles constrain cellular processes there is still room for improving the theoretical and empirical basis for estimation of some parameters. How the values of the uncertain parameters influence community structure is partly addressed by the analytical analyses in Sec. 4: the upper and lower cell sizes, the limiting levels of resources and the sizes which are most competitive for resources, and the overall biomass of the community. For example, the levels of DOC and nutrients depend inversely on the diffusive affinity *α*_D_ but increase with the coefficient of passive losses *c*_passive_. Likewise, the minimum size increase with respiratory losses while the maximum size decreases. Finally, the sizes where the dominant strategies change depend upon the affinities (Table 3). For example, increasing the light affinity coefficient *α*_L_ will decrease the sizes where there is a switch from osmotrophy to light-limited phototrophy and from light- to nutrient-limited phototrophy; increasing the clearance rate for phagotrophy *α*_F_ will decrease the size where mixotrophy becomes dominant.

### 6.2 Size-, trait-, and functional-group based plankton modelling

The central role of cell size for all vital processes makes it an obvious choice as the structuring variable for plankton community modelling. The community structure is then described as the “size spectrum” which generally follows the flat Sheldon spectrum. The minimal size-based model takes this approach further as also the structure of the trophic strategies with cell size – on the continuum of osmo-, photo-, and heterotrophy (Andersen et al., 2015) – emerges as a second dimension of community structure. This idea was previously based upon simple scaling arguments with only a single scaling exponent for each process (Andersen et al., 2016; Ward and Follows, 2016). Here we show that the same results holds even with the more complex scaling two-regime scaling laws for nutrient and light uptake. However, the transitions between the dominant resource uptake power laws are still responsible for the structure of the trophic strategies in the full dynamic model (Figs. 4.5b and 9b). The advantage of the minimal size-based description is that the entire community, from the smallest bacteria to the largest heterotrophic cells, are captured with one set of parameters that is universal across geography and time. The universal properties makes the model well suited for global simulations (Ward and Follows, 2016) under global change. The obvious disadvantage is of course that biodiversity is only described by cell size and the dominant trophic strategy.

Additional diversity can be introduced by adding other traits in addition to cell size. The size-based approach is closely related to a trait-based description of plankton (Kiørboe et al., 2018) (also referred to as the approach of “infinite diversity” (Bruggeman and Kooijman, 2007)). Size-based models are essentially the simplest form of trait-based plankton models where the only trait is cell size. The trait-based approach represents plankton by a select few traits that together best represent the functional diversity of plankton.

Traits are often related to investment in two competing resource uptakes or metabolic functions (Andersen et al., 2015): light harvesting vs. maximum synthesis rate (Shuter, 1979; Serra-Pompei et al., 2019), light harvesting vs. nutrient uptake (Bruggeman and Kooijman, 2007), adaptation between osmo-heterotropy and phototrophy (Bruggeman, 2009; Ward et al., 2011), between nutrient uptake, light harvesting, and phagotrophy (Berge et al., 2017). The trait-distribution of these traits are often Gaussian (normal distributed) and can be well represented simply by their optimal trait value (Shuter, 1979; Chakraborty et al., 2020), or by their moments (Wirtz and Eckhardt, 1996; Norberg et al., 2001; Bruggeman and Kooijman, 2007). Considering resource uptake traits in isolation represents a limited aspect of plankton diversity because the big variation in resource uptake parameters related with size is not represented. A full representation can be obtained by combining resource uptake traits with cell size (Terseleer et al., 2014; Chakraborty et al., 2017; Serra-Pompei et al., 2019; Chakraborty et al., 2020). The size spectrum itself, however, is continuous as shown in the analytical derivation of the Sheldon size spectrum. Descriptions where the size spectrum is only reduced to its moment or the optimal size (e.g. Acevedo-Trejos et al., 2018) may represent the changes of one group of plankton, but they are insufficient to resolve the entire community. Trait-size models therefore need to combine a full resolution of the size-spectrum (as done here) but can use optimization or moment-close to reduce the number of state variables for other traits.

Plankton diversity is traditionally represented by dividing phyto- and zooplankton into functional groups, including picoplankton, diatoms, flagellates, ciliates, etc. (Fasham et al., 1990). Parameters in each group can be calibrated to represent the dominant group in a given study area to achieve good fits with observations of the different taxonomic groups. Their power comes at the expense of introducing additional parameters and by requiring re-calibration if there are changes in the dominant species groups due to environmental change. A good example of a minimal functional-type model is the plankton model of bacteria, auto- and heterotrophic flagellates, diatom, and copepods (Thingstad et al., 2007). With the same complexity in terms of parameters as the minimal size-based model, the Thingstad model provides an explicit taxonomic resolution that is lacking in size-based models, though, of course, without the resolution of cell size. Size-based models are not replacements of functional-group type models, but the two types of models should be considered as complementary descriptions of the same system. Therefore global plankton models increasingly adopt descriptions that combine size and functional groups (Stock et al., 2014; Dutkiewicz et al., 2020, e.g.) to provide generality to functional-group type of models for global applications without inflating the parameter space, much like the combination of size- and trait-based models discussed above.

### 6.3 Additional traits related to cell size

Besides the resource harvesting traits discussed above there are other traits which relate to cell size. Here we first discuss the role of organisms that increase their physical size without increasing carbon mass (diatoms and gelatinous zooplankton), alternative forms of nutrient uptakes (diazotrophs), organisms with extreme predator-prey mass ratios (ciliates and larvaceans), the difference between bacteria and eukaryotes, and then present a suggestion for additional trait axes to represent that diversity.

Diatoms and gelatinous plankton increase their physical size by a large inert vacuole or a gelatinous body. In this way they gain the advantages of large physical size: higher nutrient uptake, higher photoharvesting rates, higher clearance rates, and lower average predation risk, without paying the cost of building and maintaining a large carbon mass. In a sense their success hinges on lowering their effective body density. The advantages of a lower body density follow directly from the size based relations developed here (Hansen and Visser, 2019), however, variable density is not explicitly represented in the model developed herein. Representing these life forms requires an additional trait, e.g. vacuole size (Terseleer et al., 2014; Cadier et al., 2020) or body density.

Diazotrophy is a dominant trophic strategy that is not represented in the minimal size-based model. Diazotrophs fix dissolved nitrogen gas and thereby break away from the diffusion limitation on uptake of bio-available nitrogen. However, they are also limited by diffusive uptakes of dissolved phosphorous and iron. Diazotrophy requires an oxygen-free environment, which forces the cell to limit the diffusion of oxygen into the cell. As the diffusion of oxygen into the cell follows the same size scaling as diffusive uptakes small cells will have a high influx of oxygen. It is therefore challenging for small cells to develop diazotrophy. While the limitations of cell size on diazotrophy have not been described in the literature the fundamental understanding of diazotrophy and the role of oxygen is available (e.g. Inomura et al., 2017). With such a description, diazotrophy could be added directly as an additional process into the minimal size-based model, without even adding a new trait dimension, and make diazotrophy an additional emergent trophic strategy.

The minimal model assumes that all cells have the same preferred predator-prey mass. Some organisms, however, may have very low predator-prey mass ratios, notably dinoflagellates (Kiørboe, 2008), while others have high ratios, notably larvaceans. The variation in preferred predator-prey mass ratios is accommodated to some degree by using a prey size preference with a wide size range *σ >* 1. However, that solution poorly resolves the importance of organisms with large predator-prey masses in oligotrophic situations where they act to transfer carbon from the dominant picoplankton towards larger body sizes with a re accessible to higher trophic levels. Not resolving higher predator-prey masses will underestimate the trophic efficiency in oligotrophic situations.

Finally, the insistence on just one governing set of parameters for all sizes ignores the difference between bacteria and eukaryotes. Bacteria have a different cell wall structure which most likely limits their functional cell mass (the *ν* factor) less than in the description developed here (Kempes et al., 2016).

Some of the limitations of the pure size-based approach can be addressed by including additional traits as other axes of diversity. We consider two additional axes to be prime candidates: vacuoles and a fast-slow life history axis. The vacuoles represent organisms with a lower density (diatoms) and the methodology has been successfully developed previously (Terseleer et al., 2014; Cadier et al., 2020). Technically, vacuoles are introduced as an additional size-spectrum with either a fixed vacuole size or a vacuole size which is optimized dynamically. The other trait axis would be a representation of a slow-fast life history continuum. This axis would represent how some species invest in high clearance rates and high maximum synthesis rates to achieve a fast dominance in high resource environments, while other invest in high competitive ability – low limiting resource and low respiration – and/or defence to lower the predation risk. These investments comes with trade-offs. The trade-offs between investments in resource harvesting and synthesis is somewhat understood (Andersen et al., 2015) (but see Kiørboe and Thomas, 2020), however, the investments in defence are more subtle. Recent developments in understanding the trade-offs between clearance rates and predation risk of flagellates from direct fluid mechanical simulations provides a first-principle avenue to parameterize this crucial trade-off (Nielsen and Kiørboe, 2021). Incorporating the fast-slow life history axis would also address some of the scatter in the purely size-based data of clearance rate and maximum synthesis rate (Figs. 4 and 5).

### 6.4 Conclusion

Despite the primitive representation of plankton diversity, the minimal size-based model forms a backbone on which to add other complications. Its strength is conceptual simplicity and a small set of universal parameters tied to first principles. The main effects observed in the minimal model will also be manifest in more complex size-based models, and as such the model is a useful tool to understand the mechanics of more complex size-based models. The importance of the additional complications – vacuoles, diazotrophy, high predator-prey mass ratios, or other functional groups – can be assessed with reference to the minimal size-based model. While the model is not intended as an operational biogeochemical model, the computational simplicity of the minimal model makes it useful as a basis for further theoretical ecological insights.

## Code

R code to generate all figures on github: https://github.com/Kenhasteandersen/FirstPrinciplesPlankton.

The code also includes a web-based simulator, which can be found on: http://oceanlife.dtuaqua.dk/Plankton/R.

## Acknowledgements

Thanks to all the people who helped test and validate the models: Anton Almgren, Mathilde Cadier, Lisa Eckford-Soper, Trine Hansen, Amalia Papapostolou, Camila Serra-Pompei, and Yixin Zhao. Thanks to Saeed Asadzadeh for discussions about the scaling of clearance rate with size, and to Ben Ward and an anynymous reviewer for countless constructive comments. This work was supported by the VKR Center of Excellence Ocean Life, the Gordon and Betty Moore foundation, the Simons’ foundation grant No. 931976, and the European Union’s Horizon 2020 research and innovation program under grant agreement No. 869383 (ECOTIP).

## A Calculation of effective prey preference for discrete size groups

The effective prey preferences function between size groups of predators *i* and prey *j* is calculated by integrating over the prey size preference (Eq. 30). The encountered prey in size group *j* by all predators in group *i* is:

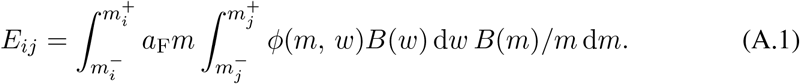

Here, *B*(*m*) represents the normalized biomass spectrum. We assume a Sheldon distribution, i.e., *B*(*m*) *∝ m^−^*^1^. With the discrete prey and predator groups we write the encountered food as:

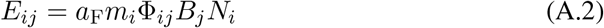

Where *B_j_* is the total biomass in group *j*, *B_j_* = *B*(*w*) d*w* and *N_i_* is the total abundance of predators *N_i_* = *B*(*m*)/*m* d*m*. Equating the two terms and isolating Φ_*ij*_ gives:

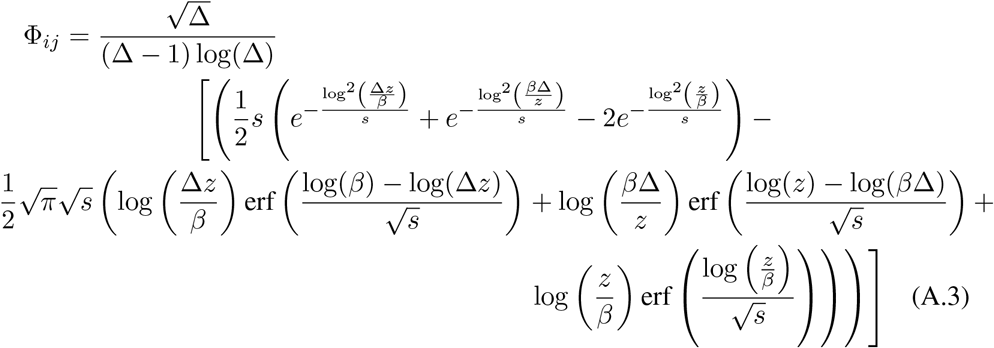

where *s* = 2*σ*^2^ and *z* = *m_i_/m_j_* and Δ = *m*^+^/*m^−^*.

---

**Figure A.1:**
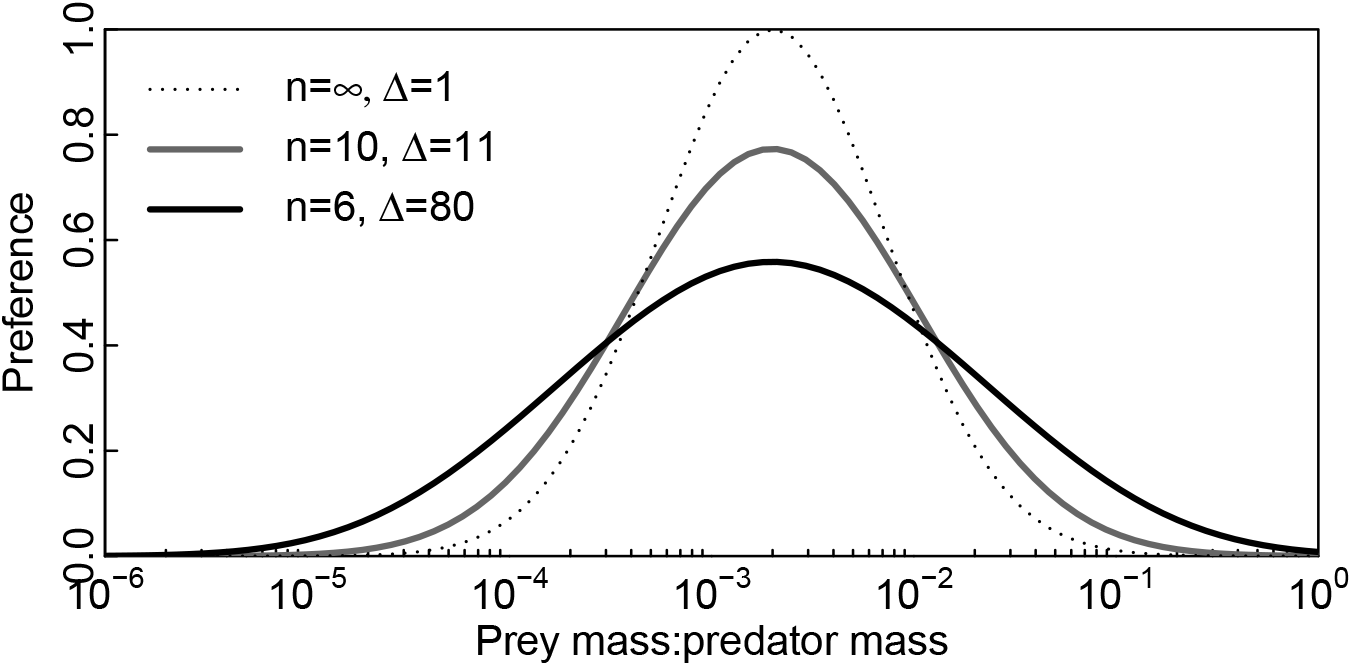
Size preference function for different grid expansions (Δ) and number of size groups (*n*).

